# Histone variant H2B.Z acetylation is necessary for maintenance of *Toxoplasma gondii* biological fitness

**DOI:** 10.1101/2023.02.14.528480

**Authors:** Laura Vanagas, Daniela Muñoz, Constanza Cristaldi, Agustina Ganuza, Rosario Nájera, Mabel C. Bonardi, Valeria R. Turowski, Fanny Guzman, Bin Deng, Kami Kim, William J. Sullivan, Sergio O. Angel

**Author notes:** Corresponding authors: Sergio O. Angel. Laboratorio de Parasitología Molecular, IIB-INTECH, CONICET-UNSAM, Av. Intendente Marino Km. 8.2, C.C 164, (B7130IIWA), Chascomús, Prov. Buenos Aires, Argentina., E-mail address (SO Angel) Laura Vanagas. Laboratorio de Parasitología Molecular, IIB-INTECH, CONICET-UNSAM, Av. Intendente Marino Km. 8.2, C.C 164, (B7130IIWA), Chascomús, Prov. Buenos Aires, Argentina., E-mail address (L Vanagas).

## Abstract

Through regulation of DNA packaging, histone proteins are fundamental to a wide array of biological processes. A variety of post-translational modifications (PTMs), including acetylation, constitute a proposed histone code that is interpreted by “reader” proteins to modulate chromatin structure. Canonical histones can be replaced with variant versions that add an additional layer of regulatory complexity. The protozoan parasite *Toxoplasma gondii* is unique among eukaryotes in possessing a novel variant of H2B designated H2B.Z. The combination of PTMs and the use of histone variants is important for gene regulation in *T. gondii,* offering new targets for drug development. In this work, *T. gondii* parasites were generated in which the 5 N-terminal acetylatable lysines in H2B.Z were mutated to either alanine (c-Myc-A) or arginine (c-Myc-R). c-Myc-A mutant only displayed a mild effect in its ability to kill mice. c-Myc-R mutant presented an impaired ability to grow and an increase in differentiation to latent bradyzoites. This mutant line was also more sensitive to DNA damage, displayed no virulence in mice, and provided protective immunity against future infection. While nucleosome composition was unaltered, key genes were abnormally expressed during *in vitro* bradyzoite differentiation. Our results show that the N-terminal positive charge patch of H2B.Z is important for these procceses. Pull down assays with acetylated N-terminal H2B.Z peptide and unacetylated one retrieved common and differential interactors. Acetylated peptide pulled down proteins associated with chromosome maintenance/segregation and cell cycle, opening the question of a possible link between H2B.Z acetylation status and mitosis.

## Introduction

Histones proteins function to form nucleosomes that regulate packaging of DNA, thereby affecting DNA transcription, replication, and repair. In addition to the four canonical histones (H2A, H2B, H3 and H4), variant histones exist that can be substituted into the nucleosome, affecting its properties. Canonical and variant histones are also subject to a wide array of post-translational modifications (PTMs) that can alter the nucleosome or recruit chromatin remodeling machinery. The combination of PTMs and exchange of histone variants is believed to be important for gene regulation in the protozoan parasite *Toxoplasma gondii,* offering new targets for drug development [1–3].

*T. gondii* is a member of phylum Apicomplexa and infects between 10-90% of the population depending on the country [4], presumably influenced by dietary habits and environmental conditions [5]. *T. gondii* converts from rapidly growing tachyzoites, which cause acute infection, to slow or non-growing bradyzoites, which cause chronic infection and typically reside in the brain [6]. Although *T. gondii* infection is asymptomatic in healthy people, untreated clinical toxoplasmosis can be lethal in immunocompromised individuals such as transplant or HIV patients, and can produce severe disease during congenital infection, especially in early stages of development [7, 8]. Chronic toxoplasmosis has also been linked to brain tumor predisposition, attention-deficit/hyperactivity disorder, obsessive-compulsive disorder and schizophrenia [9–13].

The *T. gondii* life cycle is comprised of sexual and asexual stages. While sexual replication takes place only in felids, *T. gondii* is able to replicate asexually as tachyzoites in any nucleated cell in any warm-blooded vertebrate. Following infection, tachyzoites convert into bradyzoites, which are house in tissue cysts that are impervious to immunity and current drug treatments. Recently, a myb-like transcription factor (BFD1) was described to be necessary and sufficient to induce bradyzoite differentiation [14]. How BFD1 is recruited to stage-specific promoters remains to be determined, but likely involves interplay with chromatin remodelers and epigenetic processes that include histone PTMs and/or variant histone exchange [15]. In support of this idea, treatment of tachyzoites with inhibitors of histone deacetylase 3 (HDAC3) initiates bradyzoite differentiation [16]. The roles of histone variants in *T. gondii* remain poorly understood.

*T. gondii* expresses histone variants that we have previously shown mark functional regions of the genome [17]. H2A.Z is a ubiquitous variant that has been implicated in both transcriptional activation and gene silencing in eukaryotes, and in fine regulation of important processes as DNA damage repair [18–20], which is also present in *T. gondii* [21]. Unlike H2A.Z, only specialized isoforms of the H2B histone family have been described in nature, except for apicomplexan parasites [22–25] and trypanosomatids [26], in which a H2B.Z and a H2Bv variants, were identified, respectively.

Histones H2A.Z and H2B.Z form a dimer that localizes with the transcriptional activation mark H3K4me3 in promoter/transcriptional start site (TSS) regions surrounding the nucleosome-free region upstream of the transcription start site [17]. In addition, H2B.Z and H2A.Z histones localize to the gene bodies of silent genes, including repressed stage-specific genes, suggesting a role in the regulation of stage transitions. In *T. gondii*, both H2A.Z (TGGT1_300200) and H2B.Z (TGGT1_209910) are essential during the lytic cycle with CRISPR fitness scores of −5.08 and −4.05, respectively [27]. By mass spectrometry, both histone variants were found to be hyperacetylated at the N-terminal domain, whereas few or no acetylation marks were identified on canonical H2A, H2B and H2A.X [28]. *T. gondii* H2B.Z was shown to be acetylated at 5 lysine residues in its N-terminal tail. The H2A.Z N-terminal tail has 10 acetylatable lysines, in which lysine 18 can also be methylated [28]. H2A.Z N-terminal tail acetylation has been widely shown to be a hallmark for active chromatin whereas N-terminal methylation is associated with gene silencing [29–32], suggesting it is essential for changes in gene expression during cell differentiation [33–37]. While the role of N-terminal lysine acetylation in H2B.Z has not yet been studied, N-terminal acetylated canonical H2B is associated with some, but not all, active genes in vertebrates [38]. Interestingly, H2A.Z in most eukaryotes only bears 4-5 acetylatable lysine residues in the N-terminus, while in protists like *T. gondii, Plasmodium* or *Tetrahymena termophila,* this number is considerably higher (between 5 and 16). Moreover, the presence of a *T. gondii* double variant nucleosome that altogether carries 15 acetylatable lysines that could be regulated is highly intriguing.

In the present work we studied the role of the five *T. gondii* H2B.Z N-terminal tail lysines, using a mutagenesis strategy; we also pursued a gene knockout strategy for H2B.Z. In an RH strain background, we generated *T. gondii* that possess mutated versions of H2B.Z: c-Myc-R, in which the five N-terminal acetylatable lysines were replaced by arginines, and c-Myc-A, in which they were replaced by alanines. These lines were analyzed for their ability to grow *in vitro*, differentiate to bradyzoites *in vitro* and produce virulence in mice. In addition, expression of key genes was studied, as well as nucleosome composition and DNA damage sensitivity. Our results suggest that regulation of the N-terminal positive charge patch by lysine acetylation, rather than the histone code, is essential for the aforementioned biological processes. We have also found some differential interactors for an acetylated N-terminal H2B.Z peptide, compared to a unacetylated one, opening new questions on the function of this modification and this histone variant. The implications of these data on the putative role of H2B.Z is discussed.

## Results

### Sequence alignment of Apicomplexa H2B.Z N-tail

H2B.Z is a variant histone of the H2B family that appears early in the phylum Apicomplexa. Previously, it was observed that *T. gondii* and *Plasmodium falciparum* H2B.Z N-terminal tails contain five acetylated lysines (K4,K9,K13,K14 and K18) [28, 39, 40]. The conservation of the five acetylatable lysines is clear from the alignment of H2B.Z N-terminal tail sequences from different Apicomplexa (Fig. 1A). This profuse number of modifications in the N-terminal tail, concomitant with the nucleosome partner, H2A.Z, in which several acetylatable lysines are present, make this double variant nucleosome a likely feature for chromatin modification relevant to gene expression, replication and/or DNA damage responses.

**Figure 1.**
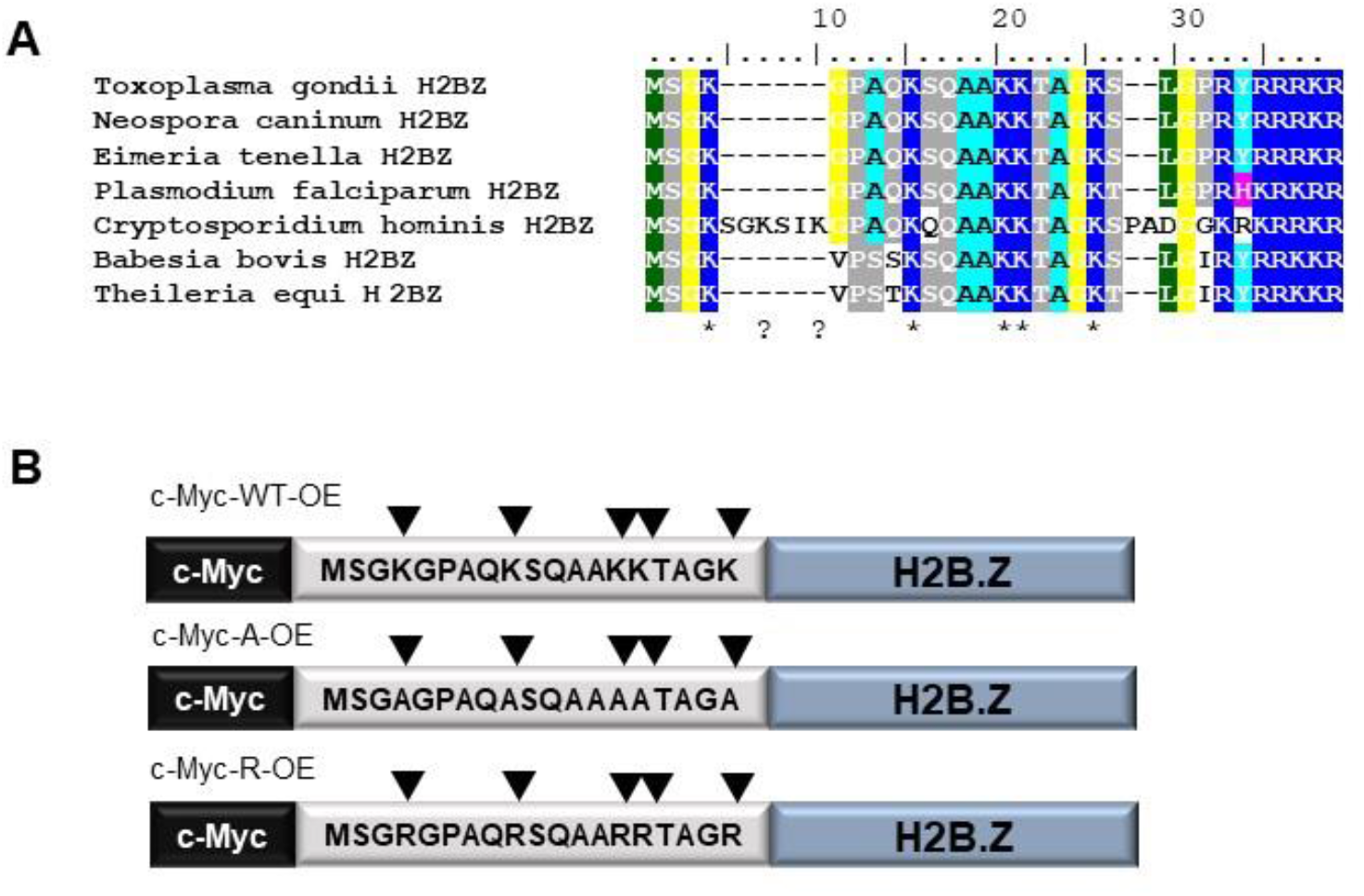
H2B.Z N-terminal tail acetylation and mutagenesis strategy. **A.** Representative Apicomplexa H2B.Z N-terminal tail alignment. *Toxoplasma gondii* (TGME49_209910), *Neospora caninum* (NCLIV_004160), *Eimeria tenella* (ETH_00030960), *Plasmodium falciparum* (PF3D7_0714000.1), *Cryptosporidium hominis* (cgd7_1700-RA), *Babesia bovis* (BBOV_IV006840) and *Theileria equi* (BEWA_028110) H2B.Z protein sequences were aligned by Clustal W. The N-terminal tail region is shown. **B.** Scheme of the constructions generated in RHΔ*hxgprt* strain tachyzoites. The N-terminal region is shown with the aminoacidic sequence, with or without mutations. All constructions were performed associated to a tubulin promoter, in frame with a c-Myc tag in the N-terminal.

### Overexpression of tagged wild type and mutant versions of H2B.Z and Knock out of endogenous *H2B.Z*

Attempts to obtain a mutant cell line by deletion of the *h2b.z* gene were unsuccessful (data not shown). The failure to obtain an H2B.Z. knockout is consistent with its CRISPR fitness score of −4.05, strongly suggesting it is essential [27] (ToxoDB, TGGT1_209910).

Given the high number of modifiable lysines in the H2B.Z variant unique to apicomplexan parasites, we analyzed their role by using an over-expression strategy of myc-tagged wild-type or mutant forms. We generated parasites over-expressing wild-type H2B.Z (c-Myc-WT-OE) or a mutated form in which all the acetylatable lysines in the N-terminal tail were changed to alanine, which mimics how acetylation ablates the positive charge of the lysine residues. We also over-expressed a mutated version in which the lysines were replaced with arginines, which would generate a constitutive positive charge patch but no longer support acetylation (Fig. 1B). Neither mutant form would be acetylated in this region, therefore would be incapable of interacting with bromodomain containing proteins. Correct localization of each c-Myc-H2B.Z protein was checked by immunofluorescence assay (IFA) (Fig. S1A) and by western blotting, observing two bands in every over-expressing line (Fig. S1B). To note, c-Myc-WT-OE was already used in a ChIP seq screen, showing the same genomic localization as endogenous H2B.Z when the experiments were performed using α-H2B.Z antibody [17].

After that, endogenous *H2B.Z* gene was deleted by using CRISPR methodology in the previously generated *T. gondii* lines overexpressing mutant H2B.Z versions (Fig 2A). We took advantage of the number of mutations in the N-terminal tail of the tagged version to design sgRNA specific for endogenous *H2B.Z* gene. By this strategy, we obtained *T. gondii* RH lines only expressing c-Myc-tagged H2B.Z with N-terminal mutations, which were named c-Myc-A and c-Myc-R (Fig. 2A-C, Fig. S1C-D). Each form of c-Myc H2B.Z in these parasites localized properly to the nucleus (Fig. 2B). In both CRISPR manipulated clones, a single band corresponding to c-Myc-tagged mutated H2B.Z can be observed by western blot with anti-H2B.Z and anti-c-Myc antibodies (Fig. 2C). Quantification of the band corresponding to c-Myc H2B.Z relativized to Sag1 as a charge control, was 1.43 ± 0.77 and 1.65 ± 0.82 for c-Myc-R and c-Myc-A, respectively, indicating comparable levels of expression between both mutants.

**Figure 2.**
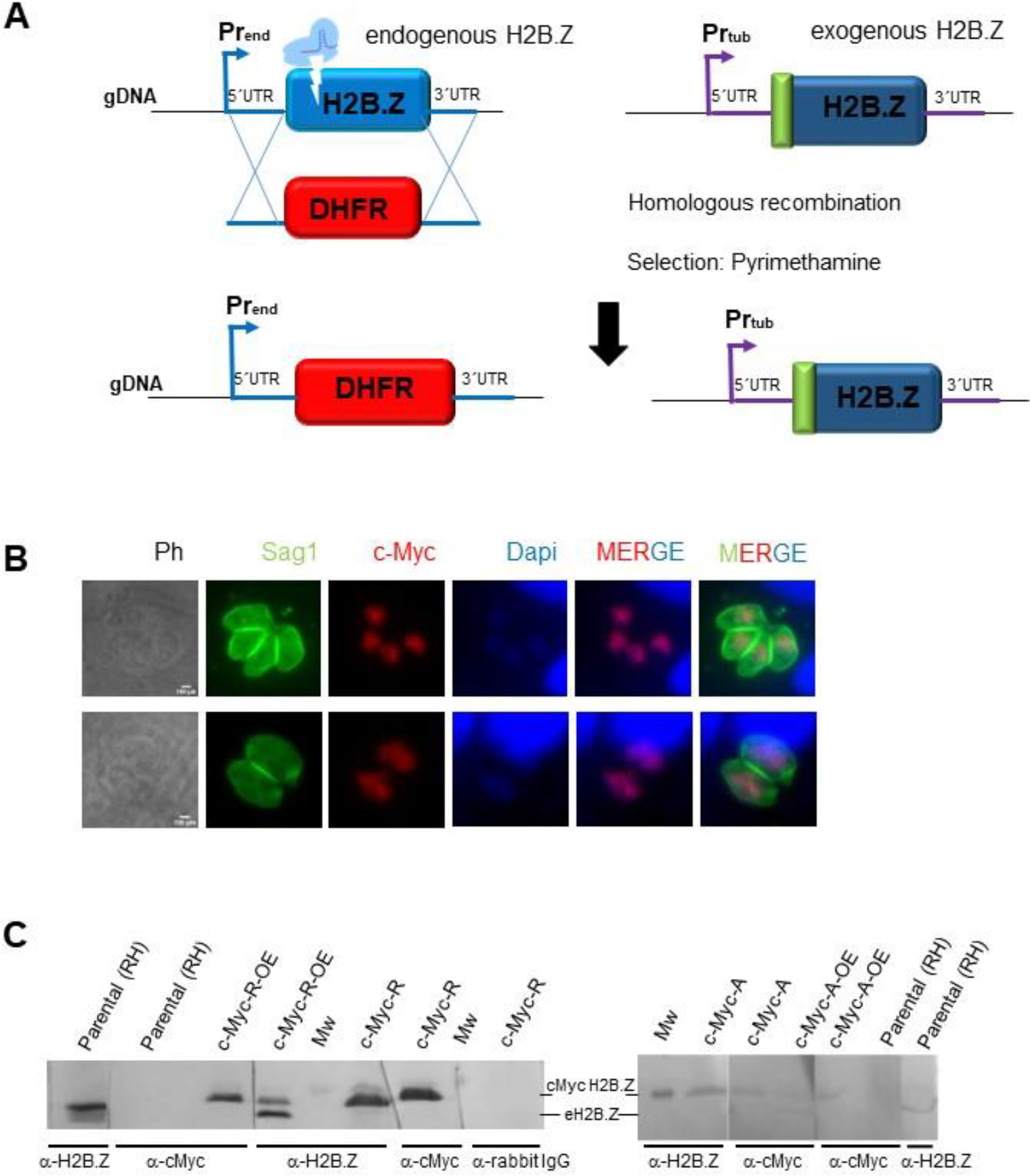
H2B.Z gene deletion strategy. **A.** CRISPR/Cas9 deletion of endogenous *h2b.z*, **s**trategy. Single guide RNA was directed to the N-terminal region of H2B.Z. Tachyzoites of one clone c-Myc-R-OE and c-Myc-A-OE were co-transfected with Crispr plasmid and a PCR product obtained with primers to amplify DHFR selection cassette from pUPRT_DHFR plasmid with homology arms (20 pb) to 5’and 3’UTR of *h2b.z*. The scheme shows the result of selection with Pyrimethamine and cloning, where parasites obtained replaced *h2b.z* by DHFR cassette, and c-Myc-H2B.Z construction (with the corresponding mutations) remained. gDNA: genomic DNA; Pr_end_: endogenous promoter; Pr_tub_: tubulin promoter. **B.** Immunofluorescence assay: coverslips seeded with Htert cells at confluency were infected with 1 tachyzoite per cell of each obtained clone and fixed after 18-24 hours. IFA was performed with α-cMyc (abCam), and α-Sag1 (mouse). α-IgG-mouse Alexa 488, and α-IgG rabbit Alexa 564 were used 1:4000 for 30 minutes. DAPI was used to stain nuclei. Scale bar=5 μm. **C.** Western blot assay. Clones confirmed by PCR were grown and lysed to run 0.5-1 x10^7^ parasites per lane, transferred to PVDF and assayed with indicated antibodies: α-c-Myc, α-H2B.Z, secondary α-rabbit IgG (negative control). Over-expressing clones (c-Myc-R-OE and c-Myc-A-OE) used for transfection, as well as the parental (RH) were loaded as controls. Mw: molecular weight marker (17 kDa band). eH2B.Z: endogenous H2B.Z.

### *In vitro* growth analysis in c-Myc-R and c-Myc-A *T. gondii* lines

We used a competitive growth assay to determine differences in parasite growth between parental wild-type and mutant parasites (c-Myc-A and c-Myc-R) obtained, which could be distinguished by the c-Myc tag. Approximately equal amounts of c-Myc positive and c-Myc negative (parental) tachyzoites were mixed and allowed to invade hTert cells, which were analyzed by IFA on different days post-infection. While the c-Myc-A parasites showed no change in growth relative to the parental line over time, the c-Myc-R line showed a marked growth deficiency compared to the parental line (Fig. 3A). As arginines conserve the positive charge but cannot be acetylated, these results indicate that the constitutive positive charge patch on the H2B.Z N-terminal tail, but not the lack of acetylation, impairs tachyzoite growth *in vitro*. Therefore, wild-type *T. gondii* must regulate the charge patch on H2B.Z N-terminal tail by PTMs such as acetylation for normal growth.

**Figure 3.**
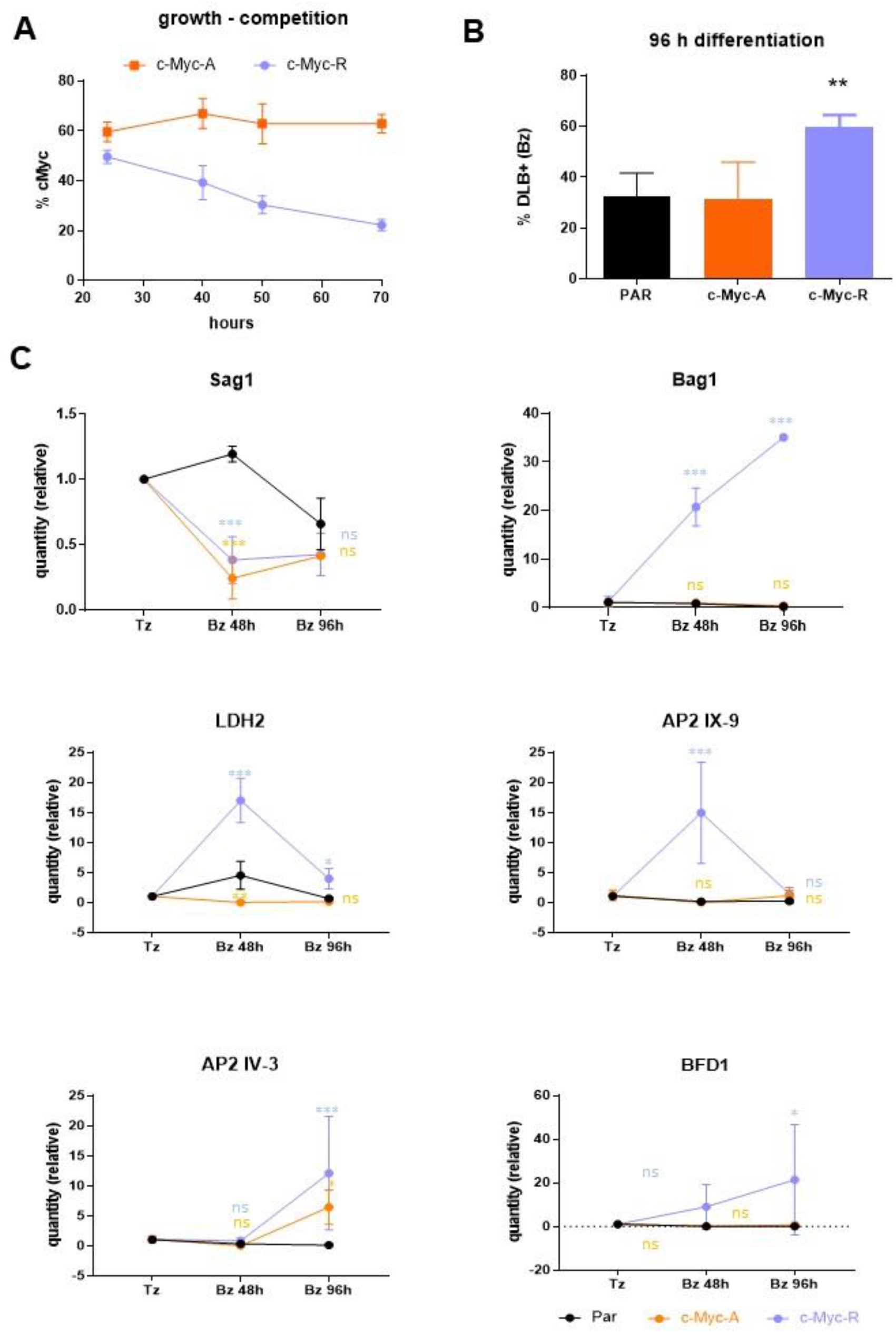
*In vitro* growth and differentiation analysis on c-Myc-A and c-Myc-R. **A.** Growth assay, performed as explained in Materials and Methods. The percentage of c-Myc positive vacuoles is expressed as relative to the total (α-Sag1), in at least 100 vacuoles per slide. Three slides per clone were counted. Representative assay of three independent experiments. **B.** *In vitro* differentiation assay was performed as explained. Percentage of DLB positive vacuoles was obtained after IFA with DLB and α-Sag1 to detect tachyzoites. Representative assay of three independent experiments. PAR: parental (RHΔ*hxgprt)*. Data was analyzed with Graphpad, and one-way Anova was performed. **: p< 0.01. **C.** Tachyzoites (Tz) of parental (RHΔ*hxgprt*); c-Myc-A or c-Myc-R were freshly collected and conserved in Trizol until processing. *In-vitro* differentiation was performed in dishes for 48 or 96 hours, and parasites were collected and conserved in Trizol. Every experiment was performed in triplicate. RNA was extracted from each sample and cDNA was obtained by reverse transcription with MMLV as explained in Materials and Methods. RT-qPCR was performed in Real time PCR equipment (Applied Biosystems) with Sybr Green reagent with sets of primers indicated in each graph, using tubulin and actin as housekeeping controls. Data was normalized to each of the housekeeping, and relative quantities to tachyzoite in each sample is plotted for each of the genes studied. GraphPad Prism was used to statisticaly analyze data, by two-way Anova. ***: p< 0.0001; **: p< 0.01; *: p< 0.05; ns: not significant.

### *In vitro* differentiation analysis in c-Myc-R and c-Myc-A *T. gondii* lines

We next studied the effect of the mutant H2B.Z variants on *in vitro* tachyzoite to bradyzoite differentiation using alkaline stress to trigger stage conversion and *Dolichos biflorus* lectin (DBL) to monitor tissue cyst wall formation. Strikingly, we found that c-Myc-R, but not c-Myc-A, showed an increase in DBL-positive vacuoles compared to the wild-type control (Fig. 3B), suggesting that the impossibility to regulate the presence of a positive charge patch in the H2B.Z N-terminal tail promotes an *in vitro* differentiation.

We also checked whether key stage-specific genes were altered in parasites expressing mutant H2B.Z. We obtained RNA from freshly lysed tachyzoites, early bradyzoites (48 hours post-differentiation), and late bradyzoites (96 hours post-differentiation) of the parental RH strain and c-Myc-A or c-Myc-R parasites. We analyzed the expression of *Sag1* (tachyzoite), *Bag1* (early bradyzoite)*, LDH2* (early bradyzoite), *AP2IX-9* (mature bradyzoites) [41], *AP2 IV-3* (in which expression is maximal at 48h post-differentiation, decreasing in mature bradyzoites) [42] and *BFD1* [14] mRNA by RT-qPCR. Actin and Tubulin were used as housekeeping genes, and data were normalized to actin. The amount of expression relative to tachyzoite in each parasite line is plotted, showing a trend of expression in c-Myc-R similar to what would be expected for a cystogenic strain with the treatment, and different from RH expression profile (Fig. 3C). Also, the expression of *BFD1* showed an increase in c-Myc-R, compared to RH (Fig. 3C). Notably, the expression profile of BFD1 in RH type I strain has not been published to date. These results are consistent with the heightened bradyzoite differentiation rate seen in c-Myc-R parasites.

### Sensitivity to DNA damaging agents in c-Myc-A and c-Myc-R lines

In other species, H2A.Z has been linked to double strand break repair driven by the non-homologous end joining (NHEJ) pathway [20, 43, 44]. Consequently, H2A.Z is associated as a responder to HU or MMS genotoxic effects [44]. We have previously characterized the anti-*T.gondii* effect of MMS, HU, and campothecin [45]. Topotecan is a genotoxic drug analog to campthotecin that produces fork collapse, and ulterior DNA double strand break repaired by HRR [46].

Given the roles of histone variants in DNA repair, we analyzed c-Myc-A and c-Myc-R parasites for sensitivity to topotecan at its IC_50_ value (20 μM, unpublished results) and MMS (50 μM) using the competitive growth assay. We observed a significative decrease in growth compared to the parental after treatment only with MMS for c-Myc-R tachyzoites (Fig. 4A). By contrast, topotecan did not produce a significant defect in growth, although c-Myc-R seems to be more sensitive (Fig. 4A). In addition, treatment with MMS produced a disorganization of vacuoles, rounded shapes and a loss of genetic material in both mutants as well as parental (Fig. 4B). *T. gondii* normally presents a synchronic replication in the parasitophorous vacuole (PV). As a result, PVs usually show 2, 4, 8, 16 and so on, tachyzoites (Tz) per PV. Treatment with MMS and Topotecan also caused a loss of synchronization, with an altered number of Tz per PV, which was also counted as “anomalous vacuole”. Anomalous vacuoles and loss of genetic material were also observed with topotecan treatment (Fig. 4B). However, quantification showed that anomalous PV are significantly more abundant in c-Myc-R than parental and c-Myc-A, with both drugs tested (Fig. 4C). Taken together, the constitutive positive charge patch generated in the c-Myc-R parasites alter the DNA damage response after MMS treatment, suggesting that acetylation status of H2B.Z N-terminal tail is likely to be modulated for proper DNA repair.

**Figure 4.**
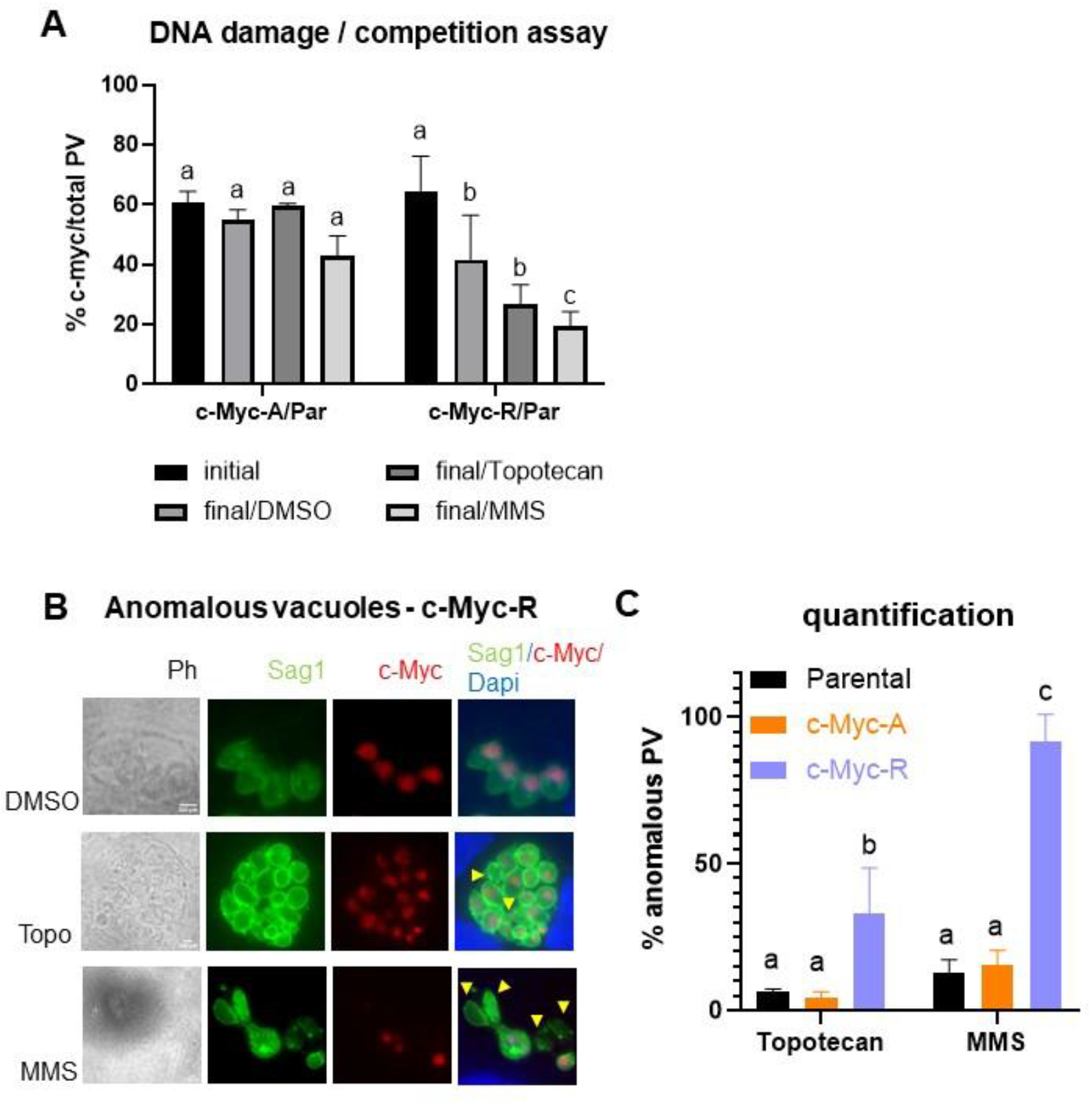
H2B.Z acetylation status is important for DNA damage sensitivity. **A.** Competition assay after DNA damage induced by Topotecan (20 μM) or MMS (50 μM). Competition assay was performed as explained, and IFA with anti-Sag1 for total vacuoles count and anti-cMyc for clones count was performed in three independent experiments in triplicate. Graph shown corresponds to one representative assay. Initial: % of c-Myc vacuoles of the total after 24 hours of infection with the mixture. Final: % of c-Myc vacuoles of the total after 72 hours of treatment with each drug or DMSO as control. c-Myc-A/Par: mixture of c-Myc-A with parental (RHΔ*hxgprt*) tachyzoites; c-Myc-R/Par: mixture of c-Myc-R with parental (RHΔ*hxgprt*) tachyzoites. GraphPad Prism 8 statistical ANOVA analysis, multiple comparison where DMSO final is compared to initial in each case and drugs data is compared to DMSO: a: ns; b: p<0.05; c: p<0.01. **B.** Vacuoles affected by drugs. Representative images of most affected parasites (c-Myc-R) with both drug treatments. DMSO images are included as control. Yellow arrowheads point to parasites with loss of genetic material. **C.** Quantification of anomalous vacuoles. Anomalous and normal vacuoles were counted in at least 50 fields randomly chosen, per clone, experiment and drug treatment. Results were plotted in GraphPad Prism 8 and statistical analysis by Multiple comparisions was performed. a: ns; b: p<0.05; c: p<0.001.

### *In vivo* analysis in c-Myc-R and c-Myc-A *T. gondii* lines

Proper gene regulation is required for tachyzoite propagation in different tissue environments and differentiation *in vivo*. Progression of tachyzoite infection of these mutants was analyzed in C57BL/6 mice by inoculating 100 tachyzoites per *T. gondii* line. As expected, parental parasites displayed high virulence, killing mice within 7 days (Fig. 5A). c-Myc-A tachyzoites were also lethal to mice, but with a delay in the time of death (Fig. 5A). According to Mantel-Cox and Gehan-Breslow-Wilcoxon tests, the survival curves are significantly different (p>0.0001), suggesting that acetylation of at least some of the 5 N-terminal tail lysines is relevant for parasite progression *in vivo*.

**Figure 5.**
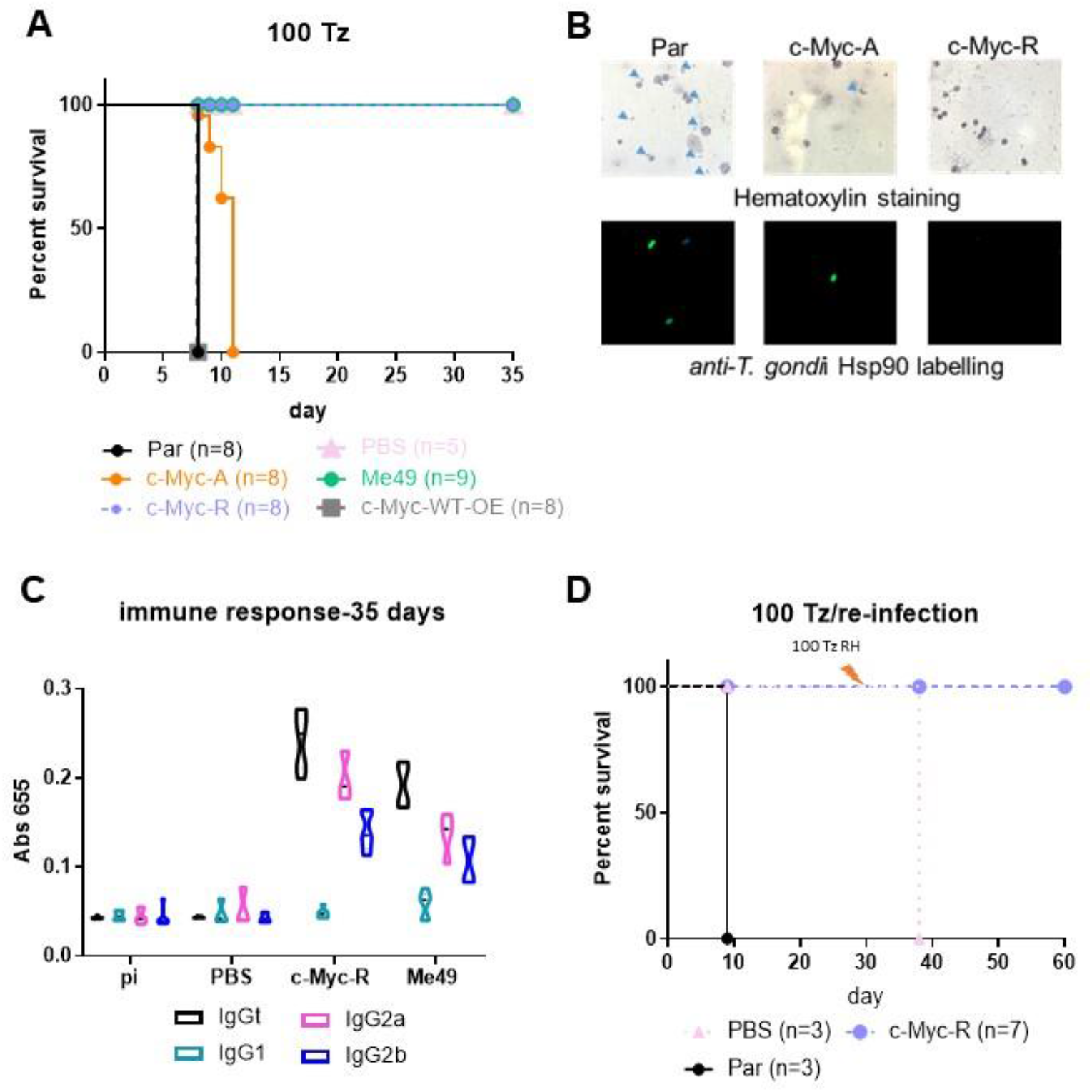
c-Myc-R tachyzoites are not virulent i*n vivo* and elicit a protective immune response. **A.** Survival assay. 10 mice C57BL/6 were intra-peritoneal infected with either 100 tachyzoites of parental (RHΔ*hxgprt*), and one clone of each KO line: c-Myc-R and c-Myc-A. As controls we also infected mice with c-Myc-WT-OE, vehicle (PBS) and with Me49 tachyzoites (type II). The survival curve shown corresponds to one assay, representative of three independent experiments. **B.** Acute infection. At day 5, 2 mice of each group were sacrificed, and intra-peritoneal liquid was extracted and analyzed in the microscope, either by hematoxylin staining (upper panel), or by IFA with α-HSP90 antibody (lower panel). Arrowheads in the upper panel show tachyzoites. **C.** Evaluation of immune response by ELISA. Blood samples were extracted from surviving mice at days 0 (Pi), 14, 21 and 35. Pooled sera from mice inoculated with PBS (negative control) or infected with Me49 (positive control) were assayed. Total IgG and subtypes (IgG2a, IgG2b and IgG1) were analyzed. In the graph only day 35 is shown. **D.** Re-infection assay. The indicated number of mice were infected with 1000 parental (Par) or c-Myc-R tachyzoites or inoculated with PBS. After 30 days, surviving mice were re-infected with 100 parental RH tachyzoites. The survival curve shown corresponds to one assay, representative of three independent experiments.

In contrast, c-Myc-R and Me49 infected mice, as well as non-infected (PBS group) survived until sacrifice on day 30-35 (Fig. 5A). c-Myc-WT-OE was used a control that the tagged version is not responsible *per se* of any affect observed. *T. gondii* line expressing both endogenous H2B.Z and c-Myc-H2B.Z-R (c-Myc-R-OE) previous to CRISPR procedure, were as lethal as the parental line (data not shown), suggesting that the randomly integrated ptub-*H2B.Z-R* gene is not responsible for the loss of virulence and that co-expression of H2B.Z wt and H2B.Z-R mutant histone did not generate a dominant negative *T. gondii* line. Infection with 1,000 and 10,000 c-Myc-R tachyzoites were also nonlethal to mice (Fig S2A).

To confirm acute infection took place, hematoxylin and IFA staining of the peritoneal fluids at day 4 post-infection were performed (Fig. 5B). Intraperitoneal tachyzoites could be detected only with parental and c-Myc-A *T. gondii* lines infections, but not with c-Myc-R parasites (Fig. 5B). Surviving animals (c-Myc-R and Me49 *T. gondii* lines) were analyzed to detect antibodies against *T. gondii*. An increase in total IgG in mice infected with c-Myc-R tachyzoites was observed, comparable to Me49, which indicated that an initial infection had taken place (Fig. 5C, Fig. S2B). The analysis of IgG subtypes showed a Th1-like humoral response in both cases, associated to IgG2a and higher for IgG2b (normal for C57/BL6 mice), typical for an intracellular parasite infection (Fig. 5C, Fig. S2B). Following sacrifice, brains of surviving mice were processed for cyst detection by direct observation with optical microscopy and DBL staining. Tissue cysts were observed in brains of mice infected with Me49 strain, but not in those infected with c-Myc-R parasites (data not shown).

To determine if infection with c-Myc-R parasites elicits an adequate immune response, mice previously infected with 1,000 c-Myc-R tachyzoites were reinfected with 100 tachyzoites of the highly virulent RH parental line. c-Myc-R infected mice survived until sacrifice, while all mice in the control group died at day 38 (day 8 after re-infection) (Fig. 5D). These results suggest that c-Myc-R tachyzoites must have been capable of invading cells and producing a stimulation of the cellular immune response adequate to control a challenge infection before clearing.

### H2B.Z acetylation status is involved in regulation of ROP proteins expression

The ability of the c-Myc-R line to establish a protective immune response suggest these parasites can invade the host cells but are efficiently controlled by the immune system. Among the *T. gondii* virulence proteins that exert protection from PV establishment are the polymorphic rhoptry proteins Rop5 and Rop18, which disrupt the γIFN-inducible IGR pathway [47–51]. Here, we used antibodies against *T. gondii* Rop5 (TGGT1_411430) and Rop18 (TGGT1_205250) recombinant proteins for western blot analysis. Rop5 was present in parental, c-Myc-A, and c-Myc-R parasites, whereas Rop18 only in the parental line (Fig. 6A). However, quantification of the band intensities showed a significative decrease in Rop5 abundance in c-Myc-R parasites and a lack of Rop18 protein in both c-Myc-A and c-Myc-R (Fig. 6B and Fig. S3A). To test whether the absence of Rop18 was a consequence of the insertion of the H2B.Z construction in that locus, we performed the same experiment using the over-expressing tachyzoites obtained prior to CRISPR manipulation. As shown in Fig.S3 (B, C) both Rop5 and Rop18 are detectable in every clone similar to parental. Co-localization analysis showed a correct localization of Rop5 in all *T. gondii* lines whereas Rop18 could only be detected in the parental line (Fig. 6C). Of note, we also detected a decrease in the fluorescence intensity of Rop5 in c-Myc-R parasites (Fig. 6D). RT-PCR analysis indicated that both genes were expressed in every *T. gondii* line (Fig. S3D), suggesting that alteration of Rop5 and Rop18 expression in the c-Myc-R line is likely at the post-transcriptional level.

**Figure 6.**
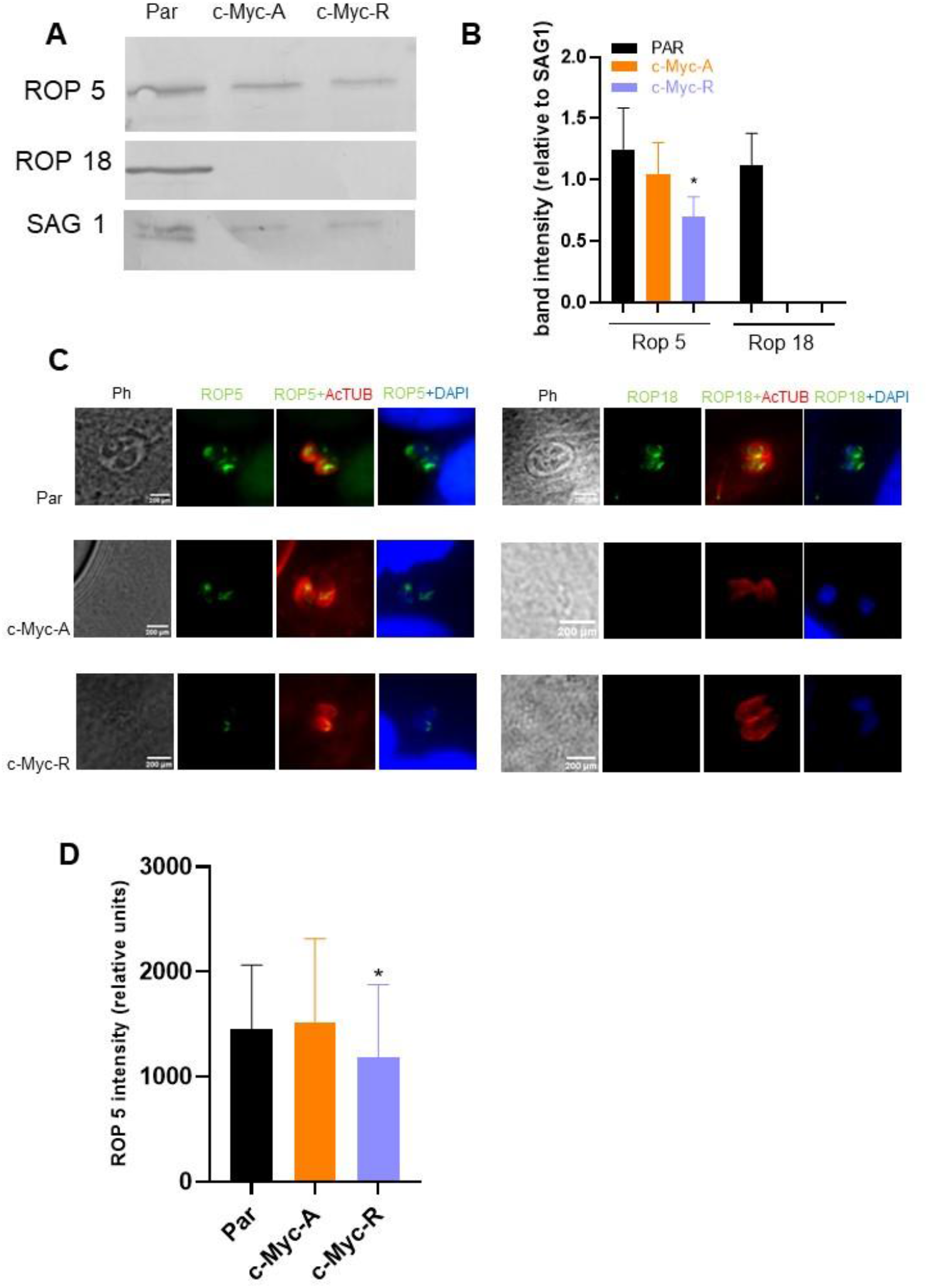
Expression of ROP proteins is affected by H2B.Z acetylation. **A.** Western blot identification of Rop5 and Rop18. The image is representative of three independent experiments, in which tachyzoite lysates of parental (RHΔ*hxgprt*), c-Myc-A or c-Myc-R were run in SDS-PAGE, transferred to PVDF membranes and assayed with anti Sag1 (charge control), anti Rop5 and anti-Rop18 antibodies. **B.** Quantification of band intensities, relative to Sag1 band intensity in each lane. Image J software was employed to quantify relative intensities and graphed with GraphPad Prism 8. Average plus SD of three independent assays is represented. *: p<0.05. **C.** Immunofluorescence assay. Htert confluent slides were infected with tachyzoites of parental (RHΔ*hxgprt*), c-Myc-A and c-Myc-R and fixed after 24 h for immunofluorescence analysis using anti Rop5 and anti Rop18 antibodies (green). AcTubulin in red, was used to stain parasites and DAPI for the nuclei. **D.** Rop5 intensity quantification. Image J software was used to quantify the antibodies intensities in three independent experiments, by triplicate, at least 10 vacuoles per slide. The graph shows the average intensity in relative units; statistical analysis was performed by GraphPad Prism 8 software. *: p<0.05.

### Double variant nucleosome and transcription is not altered in *T. gondii* mutant lines in normal conditions

Since H2BZ is part of the double variant nucleosome with H2AZ, which localizes at the promoter region of active genes, it was expected that expression alterations would occur at the transcriptional level rather than post-trancriptionally. To determine if the observations for Rop5 and Rop18 were a specific situation, we looked closer at the role of H2B.Z in transcription. We first analyzed the double variant (H2A.Z/H2B.Z) formation in both mutant lines by co-immunoprecipitation (co-IP) assays by using commercial anti-c-Myc agarose. Mononucleosomes of c-Myc-A and c-Myc-R lines were obtained after MNAse digestion before co-IP (Fig. 7A). In figure 7B it can be observed that both c-Myc-A and c-Myc-R H2B.Z interact with H2A.Z, confirming the assembly of the double variant nucleosome already found in *T. gondii*. We also decided to study the interaction of the double variant nucleosome with acetylated H3, a known mark of active chromatin. Both c-Myc-A and c-Myc-R H2B.Z interacted with this acetylated histone, which indicates that lack of acetylation of H2B.Z does not alter this PTM crosstalk (Fig. 7B and Fig. S4). Quantification of the IP band intensities relative to the respective input band showed that both *T. gondii* lines exhibit similar interaction levels with H2A.Z and acetylated H3 (Fig. 7C). These data indicate that the double variant nucleosome was not altered as well as its interaction with an open chromatin marker (acetylated H3).

**Figure 7.**
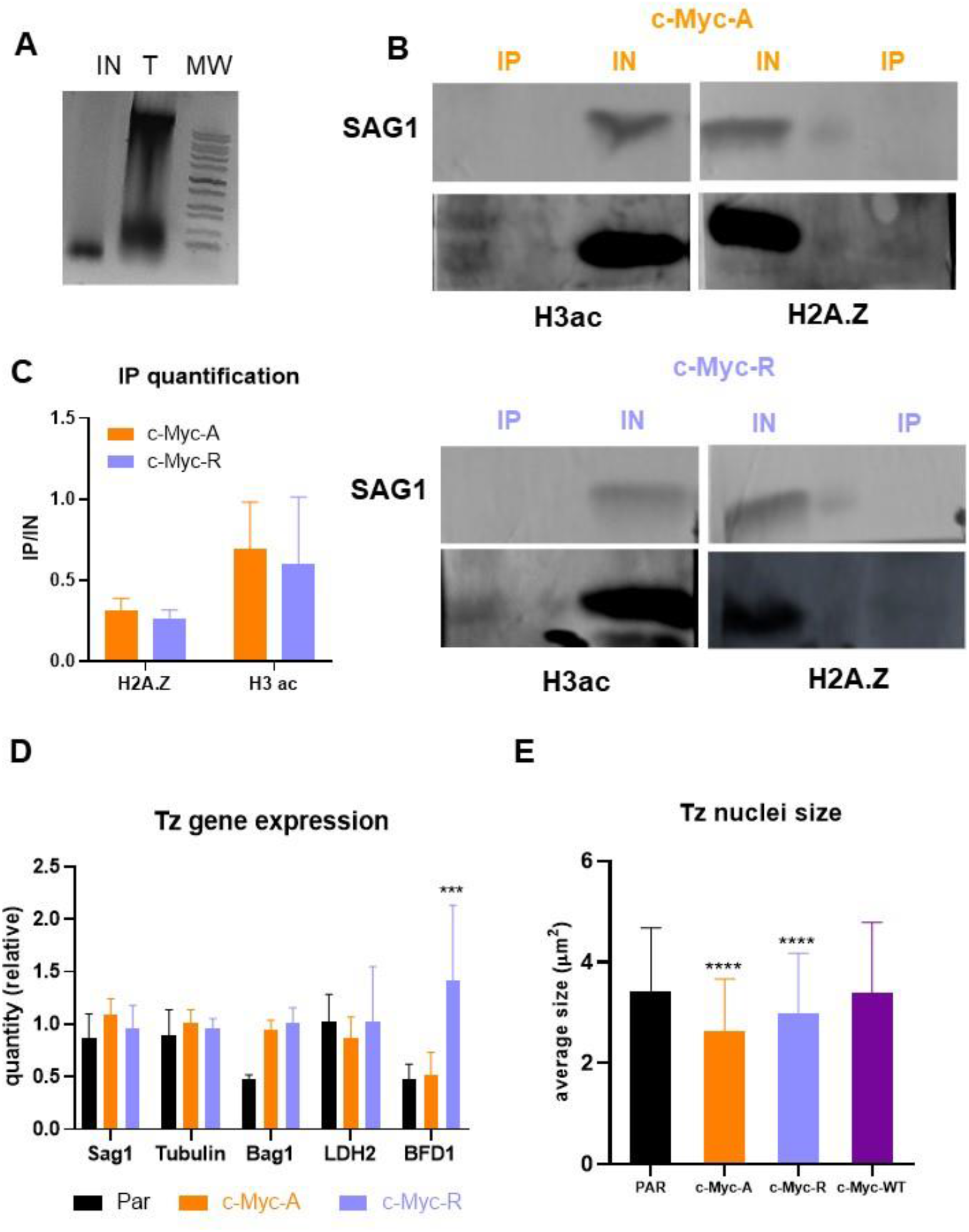
Nucleosome composition and chromatin structure is maintained in c-Myc-A and c-Myc-R. **A.** Mnase digestion of tachyzoites for co-immunoprecipitation analysis. Tachyzoites were MNase digested as explained in Materials and Methods, and an aliquot of the total tachyzoites and the digestion was precipitated and run in 1% agarose gel. A representative image of digestion is shown. **B.** Co-immunoprecipitation-WB analysis. Assays were performed in triplicate as explained in Materials and Methods. A representative image of Western-blot analysis is shown for c-Myc-A and c-Myc-R. IP: immunoprecipitation; IN: input. H3ac: western blot revealed with commercial anti H3acetylated antibody. H2A.Z: western blot revealed with anti H2A.Z antibody performed in the lab. Sag1: western blot revealed with anti Sag1 antibody. Color key: Orange for c-Myc-A; lilac for c-Myc-R. **C.** Quantification of western blot band intensities. Image J software was used to quantify intensities of H3ac and H2A.Z IP bands and relativize to respective IN band intensities. Graph shows the relative intensities of three independent experiments as the average plus SD for each antibody and clone, with the same key color as before. Graphpad Prism 8 statistical analysis showed no significant differences between clones for both H3ac and H2A.Z quantification. **D.** Tachyzoites gene expression. Tachyzoites (Tz) of parental (RHΔ*hxgprt*); c-Myc-A or c-Myc-R were freshly collected in triplicate and conserved in TriZol until processing. RNA was extracted from each sample and cDNA was obtained by reverse transcription with MMLV as explained in Materials and Methods. RT-qPCR was performed in Real time PCR equipment (Applied Biosystems) with Sybr Green reagent with sets of primers indicated in the graph, using actin as housekeeping control. Data was normalized to actin, and relative quantities to RH tachyzoites in each sample is plotted for each of the genes studied. GraphPad Prism was used to statistically analyze data, by two-way Anova. ***: p< 0.0001; ns: not significant. **E.** Tachyzoite nuclei size. Tachyzoites of parental (RHΔ*hxgprt*), c-Myc-A, c-Myc-R or c-Myc-WT-OE were allowed to invade hTert confluent slides and replicate for 20 h, and IFA was performed with α-H2B.Z antibody to detect the nuclei. Image J software was used to measure the average nuclei size of at least 100 tachyzoites per slide, in three independent experiments, and this was plotted and analyzed with Graphpad Prism 8, by two-way ANOVA. ****: p< 0.0001.

We next decided to explore if transcription was altered in tachyzoites of the mutant lines. We analyzed expression of *Sag1* (tachyzoite), *tubulin* (constitutive), *Bag1* and *LDH2* (bradyzoite) and *BFD1* (master differentiation factor). Only BFD1 mRNA expression was statistically higher in c-Myc-R that in the parental or c-Myc-A (Fig. 7D), consistent with our observation that c-Myc-R parasites are more prone to differentiate into bradyzoites.

### Chromatin compaction does not explain differences observed in *T. gondii* mutant lines

The positive charge patch on H2B.Z could be associated with a more compact chromatin structure. Since the mark of H2B.Z, either wild type or mutants, and DAPI are coincident in the diameter of the nucleus, to analyze chromatin compaction, we measured the size of the nuclei of these lines compared to the parental in IFAs stained with α-H2B.Z antibody. As shown in Fig. 7E, nuclei sizes are significantly smaller that the parental in c-Myc-R parasites in accordance with the hypothesis. However, this significant difference is also seen in c-Myc-A tachyzoites, indicating that the inability of this histone to be post-translationally modified is enough to lead to this state. The sizes of c-Myc-WT-OE parasite nuclei were similar to parental nuclei, suggesting that the c-Myc tag is not responsible for the effect observed (Fig. 7E). Taken together, acetylation of the H2B.Z N-terminal tail is required for proper control of DNA compaction status.

### H2B.Z interacting proteins

While results largely implicate the positive charge patch of H2B.Z N-terminal tail as responsible for the observed phenotypes, a role for acetylation cannot be ruled out. Acetylation may explain why the c-Myc A line showed a mild defect in virulence *in vivo* and defects in chromatin compaction. These defects could be associated with the interaction with acetylation “reader” proteins. We therefore examined if there are differential proteins interacting with H2B.Z N-terminal region depending on its acetylation state. We designed two peptides, one containing two acetylated lysines (K14 and K18) and the other unacetylated; both peptides contained a biotin tag towards the C-terminal end (Fig. 8). Peptide design was performed according to Wysocka 2006 [52]. In this work, it was recommended to perform the synthesis of peptides about 20 amino acids long, with biotin conjugated through a linker on the C-terminus for N-terminal histone peptides, and with the modification positioned close to the center of the peptide. In addition, two consecutive modifications were not recommended, explaining our choice of K14 and K18.

**Figure 8.**
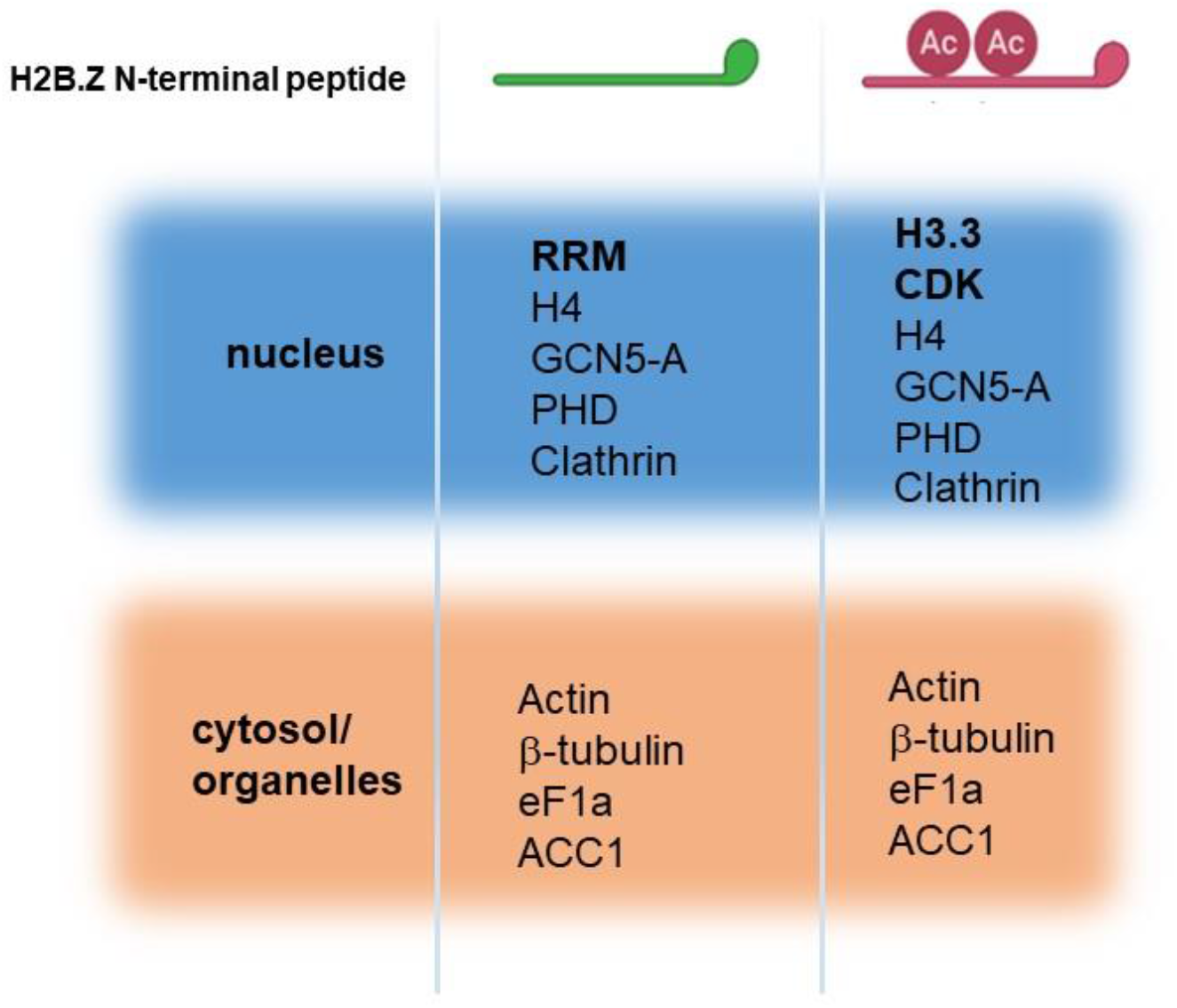
Identification of H2B.Z interacting proteins by pull-down assay. Scheme of the two different peptides synthesized is graphed above. Ac: acetylation. Most representative peptides retrieved in two independent assays for each peptide. Proteins are separated according to localization in nucleus or cytosol/organelles.

We used extracellular tachyzoites (RH) to perform the pull-down assays. After resolving on SDS-PAGE, samples were analyzed by mass spectrometry. We detected a total of 48 interactors, 18 exclusive of non-acetylated oligo, 17 exclusive of acetylated oligo and 14 present in both (Table S1). While many interactors have nuclear localization, other localizations were detected, and many typical contaminants (e.g. ribosomal subunits). Among the nuclear proteins, we can select a few that can be interacting with the H2B.Z N-terminal tail independent of acetylation status; among them, GCN5-A, histone H4 and a PHD protein (Fig. 8). Interestingly, only three nuclear proteins were differentially pulled down with strong evidence (2 independent pull downs), one of them TGME49_262620, a RRM protein (unacetylated peptide), H3.3 and a putative CDK protein (both with the acetylated peptide). Considering the nuclear genes that only appeared in 1 pull down, the acetylated peptide retrieved only genes that code for proteins associated with the maintenance and segregation of chromosomes during the cell cycle, as peptidase c50 (TGME49_262825), RecF/RecN/SMC N terminal domain-containing protein (TGME49_231170) and meiotic recombination protein DMC1 family (TGME49_216400) (Table S1). Among the retrieved peptides in both acetylated and non-acetylated pull-downs, we also detected an apicoplast localized acetyl-CoA carboxylase ACC1 (TGME49_221320) (Fig. 8 and Table S1). It is interesting to observe that an apicoplast protein displayed a significant number of peptides, indicative of a consistent interaction which could occur if H2B.Z presents in some situation an extracellular location as has been observed in other systems [53–57].

## Discussion

In *T. gondii* gene expression is associated to the cell cycle, lytic cycle and differentiation [15, 58–63]. In addition to gene expression, the presence of histone variants as well as the turnover of PTMs on histones are also associated to chromosomal organization, DNA repair or DNA replication [64]. The Apicomplexa H2A.Z/H2B.Z double variant histone nucleosome is unique and is extensively modified by PTMs including N-terminal tail acetylation. *T. gondii* epigenome has been studied, and it has been observed that this parasite uses a set of different histone modifications as a complex set of tools to control its gene expression [65]. Acetylation and methylation are associated to opposite impacts in gene expression, the first favoring expression, while the second is often a repressive mark. As an example, acetylation of lysine 31 on histone H4 (H4K31) was associated to the promoter of a nearby active gene, while in the core body of these genes H4K31me1 was detected [64]. Until now, the role of PTMs in *T. gondii* H2BZ has not been studied. However, *T. gondii* H2B.Z/H2A.Z double variant nucleosomes were found in the promoter of active genes and in the gene body of inactive genes [17]. This change in genomic location could be regulated by the different PTMs of H2B.Z and/or H2A.Z. The present work is the first attempt to decipher the role of the N-terminal acetylation of H2B.Z.

According to the results obtained in this work, the presence of a constitutive positive charge patch in the N-terminal tail of the histone (c-Myc-R mutant) causes an increase in differentiation, a decrease in growth, and an impaired DNA damage response *in vitro*. On the other hand, virulence was severely affected *in vivo* indicating some sort of regulation at least in those processes for this H2B.Z PTM. Those effects were not observed when lysine was replaced by alanine, an uncharged amino acid that mimics acetylated lysine, except a mild effect on virulence *in vivo*. For *T. termophila* when engineering a constitutively positive charge patch on H2A.Z, this was highly toxic, suggesting that its regulation has an essential function [66]. Moreover, the presence of a single acetylation site in this region was sufficient to avoid the toxic effect of the positive charge patch [66, 67]. In our case, some phenotypical alterations were observed *in vitro*, but the *T. gondii* c-Myc-R tachyzoites were viable except during mouse infection. This indicates that at least *in vitro*, if there is a toxicity due to the constitutive positive charge patch of H2B.Z, it could be partly compensated by H2A.Z, which presents 10 acetylatable lysines [28]. Together, H2A.Z/H2B.Z, would represent a “combined N-terminal tail” of 15 lysines. Studies involving the role of acetylation on H2A.Z N-terminal tail of *T. gondii* are needed to understand this.

An interesting aspect is that the *T. gondii* c-Myc-R line can grow responding to DNA damage *in vitro*, with significant but surmountable defect. This would indicate a role in DNA damage repair for this histone variant in *T.gondii*, with acetylation as a central player. However, this sensitivity was only significant for c-Myc-R tachyzoites, arguing once again in favor of the positive charge patch toxicity hypothesis in this mutant. Since H2B.Z does not exist in nature beyond Apicomplexa, we can only infer the putative role of this variant histone being part of H2A.Z/H2B.Z nucleosome. The participation of H2A.Z in DNA damage repair pathways has been studied in mammals and yeast [20, 43, 44]. In those cases, H2A.Z was found to be involved in repair due to genotoxic agents such as MMS and HU [44]. In our case, H2B.Z only demonstrated sensitivity with MMS and not with topotecan, supporting the possibility that H2B.Z/H2A.Z work together in the same function.

The contributions of effects observed *in vitro* are less difficult to ascertain in the *in vivo* model. Clearly c-Myc-R line is capable of invasion, generating an adequate immune response, but it is quickly neutralized during the infective process. This suggests that the tachyzoite is exposed *in vivo* to stronger environmental pressures than those observed *in vitro*. These pressures would include adaptation to new environments, including different cell types and organs, which may present differences in metabolism, nutrient availability, or immune responses. Under these conditions, where the tachyzoite must respond quickly, fine reprograming of gene expression and modifications in metabolic pathways would be required and the impossibility of modulating the positive charge patch of H2B.Z could hamper the ability of parasites to adapt to changing environments *in vivo*.

A possible explanation for the changes provoked by the positive charges in the N-tail could be a chromatin compaction due to interaction with the negative charges in DNA. Genome size expansion in eukaryote evolution has led to an increase in histone H2A N-terminal tail arginines. Arginine-rich histones bind more tightly to DNA, compacting chromatin and, in some cases, the nuclei are also smaller [68]. However, in this work we showed that nuclei size was smaller in both clones compared to parental and to c-Myc-WT-OE tachyzoites, indicating that the charges in the N-terminal tail are not responsible for chromatin compaction. Chromatin compaction has also been described to be regulated by linker histone H1, that stabilizes the nucleosome structure. A candidate H1 has been recently identified in *T. gondii*, and interaction of H1 with H2B.Z was observed [69]. The acetylation status of H2B.Z may be important for this interaction, leading to a more compact chromatin in both alanine and arginine mutants because of linker histone H1. Further experiments are needed to test this hypothesis.

It could be expected that the phenotypic alterations are given by defects in the formation of the nucleosome with an impact on the expression of the genes. In our analysis, both situations are not altered. Although a massive analysis of gene expression is necessary to detect specific changes, part of the study shows that the expression of Rop5 and Rop18 does not occur at the transcriptional level, but rather post-transcriptionally, affecting their translation. This regulation could be given by the presence of non-coding RNAs (ncRNAs) and there is increasing evidence for a significant role for pos-transcriptional mechanisms [70, 71]. Future studies should take this possibility into account, considering a possible role of H2B.Z acetylation, to analyze ncRNAs expression in *T. gondii*. With the results shown in this work it is only possible to correlate the decrease in one of the more virulent isoforms of Rop5 together with the absence of Rop18 protein to the loss of virulence of c-Myc-R mutant, although it cannot be ruled out that other virulence factors could be altered in these parasites. On the other hand, the absence of Rop18 protein in c-Myc-A correlates with the significative delay in mice death with c-Myc-A tachyzoites infection as reported by Resse et al [49].

We could not detect significant changes in most of the genes analyzed by RT-qPCR in normal conditions for tachyzoites, except for *BFD1* in c-Myc-R. This does not discard an association of the N-terminal tail of H2B.Z with the epigenetic modulation of gene expression in some circumstances, for example in changing environments, in an *in vivo* model. Regarding the significant expression of BFD1 mRNA, it is intriguing if it is related to the high differentiation rate observed *in vitro* for these parasites. It is important to note that the expression of the BFD1 gene is post-transcriptionally regulated [14]. Therefore, the increase in transcription *per se* is not enough to induce differentiation, but the higher levels of BFD1 mRNA could favor differentiation under alkaline stress. There is also another possibility of inducing differentiation based on epigenetic regulation. HDACi FR235222 treatment was shown to favor acetylation of H4K31ac in bradyzoite gene promoters, promoting transcription of bradyzoite genes and therefore, bradyzoite development [64]. Maybe under differentiation stress, H2B.Z is implicated in a fine regulation of gene expression through the acetylation level of its N-terminal tail. Keeping that in mind, the lack of modulation of H2B.Z in c-Myc-R could result in a different chromatin firing up the expression of bradyzoite-specific genes under alkaline stress.

Both acetylated and unacetylated peptides pull-downs share 3 nuclear proteins: GCN5A, PHD and clathrin. GCN5A was shown to impact on H2A.Z acetylation via crosstalk with H4 acetylation by means of bromodomain in p300 [72]. On the other hand, unacetylated peptide recognized one protein specifically, a RNA recognition motif (RRM)-containing protein (TGME49_262620), which was also pulled down with GCN5b acetyltransferase [60]. RRM-containing proteins are involved in post-transcriptional gene expression processes, such as mRNA and rRNA processing, RNA export, translation, localization, stability, and turnover [73]. In the case of acetylated peptide, it pulled down 2 specific proteins: a cyclin-dependent kinase regulator (CDK_ TGME49_219832) and H3.3 histone variant (TGME49_218260). CDKs phosphorylate multiple substrates that regulate the cell-division cycle in response to different cellular cues [74]. H3.3 and H2A.Z are associated to promoter regions flanking the nucleosome depleted region (NDR) of active genes [75]. Therefore, it may be likely that acetylated H2A.Z and H2B.Z would associate to this variant histone in active promoters. However, this association was not found in our genome wide analysis on tachyzoite stage [17].

In addition, CDKs have also been linked to mitosis [76, 77] and H3.3 is related to chromosome segregation, nuclear structure, and the maintenance of genome integrity [78]. In this direction, it is interesting that our pull-down assay has also retrieved, only for the acetylated peptide, other genes, all of them associated to chromosome segregation (Table S1). Among them, peptidase c50 (TGME49_262825) a separase or caspase-like protease, known to play a central role in the chromosome segregation [79]. Structural maintenance of chromosomes (SMC) proteins-like RecF/RecN/SMC N terminal domain-containing protein (TGME49_231170)-function together with other proteins in a range of chromosomal transactions, including chromosome condensation, sister-chromatid cohesion, recombination, DNA repair and epigenetic silencing of gene expression [80–82]. Finally, meiotic recombination protein DMC1 family (TGME49_216400) participate in meiotic recombination, specifically in homologous strand assimilation, which is required for the resolution of meiotic double-strand breaks [83]. These findings open an interesting question, which could implicate the role of H2A.Z/H2B.Z in the process of mitosis and chromosome segregation in the future.

It is relevant that these tachyzoites were able to generate a complete immune response, which only happens when cells are invaded by tachyzoites and not with an infection with dead or lysed tachyzoites [84–87]. In this sense, Toxovac® is a live attenuated vaccine used to prevent sheep congenital toxoplasmosis [88]. Recently, a Ca^2+^-dependent protein kinase (CDPK) knock-out in Me49 type II strain was evaluated in mice as a potential vaccine candidate [89]. The protection we observed in this work generated in mice facing an infection with virulent RH tachyzoites is encouraging to continue with the proper assays needed for the formulation of a vaccine.

## Conclusion

In this work, we have shown that N-terminal tail lysine acetylation of histone H2B.Z is important for the normal progression of *Toxoplasma gondii* life cycle. The presence of 5 positive charges in the N-terminal of H2B.Z is most likely to be detrimental for the parasite. However, this toxicity is not very high, allowing the parasite to replicate, but conferring a lower fitness to the tachyzoite. Moreover, this lower fitness is traduced in a higher sensitivity to DNA damage, a mis-regulation of gene expression conducing to a higher differentiation rate *in vitro*, and also a complete lack of virulence when facing the immune response in mice. All these effects are not explained by an heterochromatinization of DNA due to the interaction with the positive charges, because we observed that nuclei are also smaller in c-Myc-A tachyzoites. Besides, the effects cannot be explained by a dislocalization of the histone in the genome because the nucleosome is still constituted normally. However, it is quite unlikely to find a physiologically a situation where the histone would be completely unacetylated, and c-Myc-A tachyzoites may reflect the normal condition observed *in vivo*. Therefore, the similarity observed between c-Myc-A and parental tachyzoites in most experiments performed in this work, would confirm that in normal conditions H2B.Z is acetylated in *T. gondii*.

The findings in this work are not exclusively explained by regulation of gene expression. This is not surprising, as it is known that histone proteins may exert other functions, related for example to cell division. Even extra-nuclear functions are attributed to histones in many models, and research in this area could bring new insights for *T. gondii* as well. Deficiencies in *Trypanosoma cruzi* H2Bv histone variant amount present similar phenotypes as observed here, and the authors also proposed a putative extra-nuclear or extra-nucleosome function [57].

Having a better understanding of how chromatin, chromosome maintenance, DNA integrity, DNA replication, and gene expression are modulated in *T. gondii* will allow inferring about chromatin modulation in other parasites of the phylum as well as the possibility of suggesting a novel mechanism of epigenetic regulation in organisms with complex life cycles that are evolutionarily distant from fungi and vertebrates.

## Materials and methods

### Parasite culture and manipulation

RHΔ*hxgprt* strain was used in all cases and grown in standard tachyzoite conditions *in vitro*: hTERT (ATCC® CRL-4001, USA) monolayers were infected with tachyzoites and incubated with Dulbecco’s modified Eagle medium (DMEM, Invitrogen) supplemented with 1% fetal bovine serum (FBS, Internegocios S.A., Argentina) and penicillin (10,000 units/ml)-streptomycin (10 mg/ml) solution (Gibco, Argentina) at 37°C and 5% CO_2_.

### Cloning over-expression strategy

To generate the c-Myc-H2B.Z WT construct, the open reading frame (ORF) was amplified using the primers F: 5’-<u>ATGCAT</u>TCAGGGAAAGGTCCGGCACAG-3’ and R: 5’-<u>TTAATTAA</u>CTATGCACCAGAAGTCGTG-3’. For the mutant constructs, reverse primer was the same, and forward was: <u>ATGCAT</u>TCAGGG**GCA**GGTCCGGCACAG**GCA**TCTCAGGCGGCG**GCGGCG**ACCGCCGGG**GCG**TCTCTGGGAC for c-Myc-A, <u>ATGCAT</u>TCAGGG**CGC**GGTCCGGCACAG**CGC**TCTCAGGCGGCG**CGCCGC**ACCGCCGGG**CGC**TCTCTGGGAC for c-Myc-R, where lysine coding codons were replaced for alanine and arginine coding codons, respectively (in bold type). NsiI and PacI restriction sites (underlined) were engineered into the 5’ end of primers F and R, respectively, to clone into *T. gondii* expression vector PTUB8mycGFPPfTailTy-HX (a kind gift of Dr. Dominique Soldati, Université de Geneve) with a strong tubulin promoter. The resulting PCR product of the expected size was removed using a Qiaex II Gel Extraction kit (Qiagen), cloned in pGEM-T vector (Promega) and sequenced (Macrogen). The cloned ORF was digested with NsiI and PacI and ligated into the plasmid. The final plasmids were electroporated into RH strain tachyzoites lacking hypoxanthine–xanthine guanine phosphoribosyl transferase (RHΔ*hxgprt*). Transgenic parasites were selected in mycophenolic acid (25 μg/ml, Sigma) and xanthine (50 μg/ml, Sigma) and cloned by limiting dilution.

### CRISPR/Cas9 gene deletion strategy

Single guide RNAs were designed directed to the N-terminal portion of the *h2bz* gene: F: <u>AAGTT</u>GAGGCGGCGAAGAAGACCGCCG and R: <u>AAAAC</u>GGCGGTCTTCTTCGCCGCCTCA. Bsa I restriction sites were included (underlined). Both oligos were annealed and cloned into the pU6 CRISPR Universal plasmid (kindly provided by Louridós Lab, Whitehead Institute), by using the Bsa I restriction site. The plasmids were confirmed by PCR, using the gRNA as primers, were sequenced (Macrogen) and prepared for transfection in *Toxoplasma*. In order to obtain a clean KO, oligos were also designed to amplify the DHFR selection cassette from the pUPRT_DHFR plasmid available in the laboratory, with homology regions (20 bp underlined) to the 3’and 5’UTR of the *h2bz* gene (F: <u>GAAAGGTCCGGCACAGAAATCTCAGGCGGCGAAGAAGACCAAGCTTCGCCAGGCTGTAAATCC</u>, and R: <u>GAATTCCCTCTCAACATCAAATTTCACCCCGTGGTTTATTCCTCATCCTGCAAGTGCATAGAAGGA)</u>. The PCR amplification product was purified from agarose gel and used to co-transfect the parasites along with the CRISPR plasmid. c-Myc-R and c-Myc-A tachyzoites (one clone for each one was selected) were transfected and Pyrimethamine (1 μm, Sigma) was used for selection. Cas9 was detected in the nucleus of the parasites after transfection by using α-Flag antibody (Sigma, F1804). Selection was continued until three passages and cloning by limiting dilution was performed. Positive clones were selected by PCR with primers designed for that purpose: TgH2Bv-upUTR-F: GTTGTCATGCGCATTTGCATCAC; TgH2Bv-downUTR-R: GTGTGCACGCGTTATAATGAGCAC; Val-TgH2Bv-R: CATGCTCTTCTTCGACACACCAG; DHFRCX-F: GTGGCATTTCACACAGTCTCACCTC; DHFRCX-R: AGACGCAGACGCATACAACGTTAG.

### Immunofluorescence assay (IFA)

Intracellular tachyzoites grown in cover slips were fixed using cold methanol 100% for 8 min, washed with PBS and blocked with 1% BSA. Primary antibody αrH2B.Z (rabbit), αrSag1 (mouse), αrHSP90 (rabbit), αrROP5 (rabbit), αrROP18 (rabbit) produced in the laboratory and αc-Myc (rabbit, abCam ab9106 or mouse, Santa Cruz sc-42), αAcTubulin (mouse, Millipore MABT868) were incubated for 1 h at room temperature. After several washes with PBS cover slips were incubated with secondary antibodies Alexa fluor goat anti-mouse 488 and anti-rabbit 594 (Invitrogen). DAPI was used to stain nuclei. Axio Imager.M2 Microscope Carl Zeiss (Germany) with objective Plan-Apochromat 63x/1.40 Oil M27 and Video digital camera Zeiss 503 monochromatic 2.8 megapixeles, was used. Image J 1.53q (Fiji) software was used to process images.

### Western blot analysis

Tachyzoites were collected, filtered, counted and lysed by 6 cycles of rapid freezing/defreeze in hypotonic buffer, and boiled with LB for 5 minutes. 0.5 to 1×10^7^ parasites were loaded per well and resolved by 15% SDS-PAGE. Proteins were transferred to PVDF membrane for 1h at 100V. Western blot was then performed as described [90]. The primary antibodies: αrH2B.Z [22], αrH2A.Z [23], αrROP5 (rabbit) and αrROP18 (rabbit) were used at 1/5000, whereas α-c-Myc (rabbit, abCam ab9106) was used at 1/2000 and αrSag1 [87] 1/200 for 1 h at room temperature. αH3ac (rabbit, Millipore, 06-599B) 1/200, was incubated overnight. Appropriate secondary antibodies were used: phosphatase alkaline-conjugated goat anti-mouse or anti-rabbit (Sigma) along with the NBT and BCIP (Promega) detection system.

### Phenotypical assays

For growth/competition assays, coverslips seeded with Htert cells were infected with a mix of 50% parental (RHΔ*hxgprt*) and 50% of each clone. 3 coverslips per clone were infected with 0,05 to 0,1 tachyzoite per cell, incubated 10 minutes on ice, and 2 hours at 37°C in incubator for invasion. Slides were washed twice with PBS to remove tachyzoites that did not enter the cells, and fresh DMEM supplemented with 1% SFB was added. Slides were fixed at different times, up to 96 hours and IFA was performed with α-c-Myc and α-Sag1 antibodies to distinguish between parental and c-Myc positive vacuoles. Number of c-Myc positive and total vacuoles in at least 100 vacuoles, randomly choosing different fields of each slide were counted. In the case of competition assays with genotoxic drugs treatment, the experiment was modified as follows: 3 slides per clone mixture were fixed after 24 h, to stablish the initial percentage of c-Myc positive vacuoles. At this time point, media was changed in the rest of the slides for DMEM supplemented with DMSO or drugs at the concentrations indicated. These slides were fixed after 96 h, and IFA was performed and counted as explained before.

For *in vitro* differentiation assays, in a similar way using Htert confluent slides, media was changed by DMEM HEPES pH 8.1, after invasion with 1 parasite every 10 cells, accompanied with deprivation of CO_2_. Media was changed every day, and differentiation rate was estimated after 96 h by using *Dolichos Biflorus* Lectin conjugated to Fluorescein (DLB 1:200, FL-1031; Vector Laboratories) to stain the wall of the cysts in formation and α-Sag1 antibody to stain the tachyzoites. In the case of collection of parasites for RNA extraction, differentiation assay was performed in a similar way, but in t25 dishes. After 48 or 96 h parasites were forced out of the cells by 23G, 25G and 27G syringe passages.

For *in vivo* survival, virulence and differentiation assays, C57BL/6 female mice were used. For the use of animals, C.I.C.U.A.E-UNSAM 10/22 was approved. Mice were maintained in optimal conditions in the biotherium, with controlled temperature and free access to sterilized water and food. Infection was carried out intra-peritoneal in 10 mice per group with 100, 1000 or 10000 tachyzoites of parental lines or clones, depending on the assay. PBS was injected as negative control in five mice per experiment. Deceases were registered along 30-35 days in each experiment, and signs of illness were monitored daily. Two mice from each group (except PBS) were sacrificed at day 5 (when RH infected mice have symptoms of illness), and intraperitoneal fluids were collected in order to detect acute *Toxoplasma* infection. Blood samples were taken at days 0, 14, 21 and 35 and sera was frozen for ELISA assays. After sacrifice, surviving mice brains were processed for cyst observance by optical microscopy. Also, a sample was stained using DLB and checked by fluorescence microscopy.

In every phenotypical assay, three independent experiments were performed. One representative experiment is shown.

### RT-PCR and RT-qPCR

Tachyzoites or parasites exposed to 48 or 96 h differentiation stress as explained before were conserved in TriZol (Invitrogen) solution at −80°C until use. All experiments were performed in three independent replicates. RNA extraction was performed according to the manufacturer’s instructions and cDNA was obtained by means of MMLV reverse transcriptase (Promega) using oligo dT with the protocol provided with the enzyme. For RT-PCR, primers used were: SAG1: Fw: TGAGAACCCGTGGCAGGGTAA; Rv: GCTTTTTGACTCGGCTGGAA; TUB: Fw: ATGTTCCGTGGTCGCATGT; Rv: TGGGAATCCACTCAACGAAGT; ACTIN: Fw: GGGCGGTTTCATGACCTAAA; Rv: ACGTATGATGCGCGAGAAAA; ROP5: Fw: CTAGCTAGCATGGCGACGAAGCTCG; Rv: CCCAAGCTTTCAAGCGACTGAGGGCGCA; ROP18: Fw: CTAGCTAGCATGTTTTCGGTACAGCGGCCA; Rv: GCCAAGCTTTTATTCTGTGTGGAGATGTTC.

For RT-qPCR primers used were: SAG1: Fw: TGAGAACCCGTGGCAGGGTAA; Rv: GCTTTTTGACTCGGCTGGAA; BAG1: Fw:CAACGGAGCCATCGTTATCAAAGG; Rv: TAGAACGCCGTTGTCCATTG; LDH2: Fw: ACAATGGCCCAGGCATTCT; Rv: CAATAAACATATCGTGAAGCCCATA; AP2IX-9: Fw: GGGCGTTCTCAGCGTTCACT; Rv: GGCGCGTCTCATCTGTTTCA; AP2IV-3: Fw: GAGCCCATTGACCCCATGAA; Rv: GGCTTCGCTTCTTTCCGTGA; BFD1: Fw: ATGTCGGGAACGATGGTTTA; Rv: TCTTCACGGCATTCTCTGTG; TUB: Fw: ATGTTCCGTGGTCGCATGT; Rv: TGGGAATCCACTCAACGAAGT; ACTIN: Fw: GGGCGGTTTCATGACCTAAA; Rv: ACGTATGATGCGCGAGAAAA.

The primers were first assayed for efficiency using a pooled cDNA in 1 to 0.001 dilutions, and the threshold was defined for each set of primers. Melt curves were also obtained for each set. For each set of experiments, SybrGreen master solution (Roche) was used and qPCR was run in StepOne Real time equipment (Applied Biosystems). Actin and tubulin were used as housekeeping genes and data was normalized to those genes by Infostat software. For the experiments in figure 3, all sets of data were relativized to tachyzoite amplification. For experiment shown in figure 7, data from the different clones was relativized to the parental.

### ELISA assay

Blood samples were extracted from surviving mice at days 0 (Pi), 14, 21 and 35. Sera was frozen until processing for ELISA. TLA (Toxoplasma lysate antigen, obtained from fresh RH tachyzoites) in buffer Carbonate (7,13g/L NaHCO3 + 1,59 g/L Na2CO3 pH 9,5) was used to immobilize 96 well plaques (Nunc Immuno™ MicroWell™ 96 well solid plates, Sigma), o.n. at 4°C. After bocking with 5% non-fat milk in PBS-T they were incubated with sera diluted 1:100 in the same solution. Total IgG (IgGt) and subtypes (IgG1, IgG2a and IgG2b) HRP-conjugated anti-mouse, made in rat, were used 1:5000. Tetramethyl-bencidine (TMB, Invitrogen) was used to reveal, and plaques were read after 20 min reaction at 655 nm in a Synergy H1. Data was analyzed in GraphPad Prism v7. Sera from mice inoculated with PBS (negative control) or infected with Me49 (positive control) were assayed.

### Co-Immunoprecipitation (Co-IP)

Approximately 5×10^8^ RHΔ*hxgprt* tachyzoites were used for each immunoprecipitation. Tachyzoites were treated with micrococcal endonuclease (MNase) in order to obtain mononucleosomes as described [23] and Co-IP was performed as described in Dalmasso et al [23] with minor modifications. Protease inhibitor cocktail (Sigma) was added in every step. Mononucleosomes were incubated with Agarose c-Myc beads (abCam, ab1253) overnight at 4°C. Immunocomplexes were washed twice with washing buffer 1 (50 mM Tris, pH8, 200 mM NaCl and 0.05 % Igepal100), twice with washing buffer 2 (50 mM Tris, pH8, 300 mM NaCl and 0.05 % Igepal100), and twice with buffer TE (10 mM Tris, pH8, 1 mM EDTA); then resuspended in 60 µl of SDS-PAGE loading buffer. Samples were boiled for 5 min and 20 µl were loaded per well in a 15% SDS-PAGE gel for immunoblotting. The absence of contaminating proteins was corroborated by Western blot with murine anti-SAG1antibody [87]. Quantification of the bands was done with Image J 1.53q (Fiji) software.

### Tachyzoites nuclei size assay

Three slides with confluent hTert cells were infected as explained before with parental or c-Myc-A/c-Myc-R/c-Myc-WT-OE clones and let replicate for 24 hours. Slides were fixed and IFA was performed as explained before using αrH2B.Z to stain nuclei. Images were processed with Image J 1.53q (Fiji) software to measure the size of the nuclei, by delimiting them with the appropriate tool in at least 50 vacuoles in fields randomly chosen, for each slide and each independent experiment. Data was graphed and statistical analyzed with GraphPad Prism 8 software.

### Peptide synthesis and characterization

Peptides were synthesized by solid-phase multiple peptide system using tea-bags with Fmoc/tBu strategy according Guzmán [91], briefly: Rink amide resin (0.55 meq/g) was used as solid support and Fmoc amino acids, including acetylated lysine (Iris Biotech). Peptides were couplet to biotin for their detection The peptide cleavage was performed with a solution of Trifluoracetic acid/triisopropylsilane/1.2-ethandithiol /water (92.5 /2.5 /2.5/2.5), washed with cold ether dried and lyophilized. Peptides were characterized by HPLC and electrospray ionization mass spectrometry, and purified by C18 cartridges (United Chemical Technologies, Bristol, PA, USA) before use.

Peptides used in the pull-down assays were: Un-acetylated: GPAQKSQAAKKTAGKSLGPRK(Biotin) Acetylated: GPAQKSQAAK(Ac)K(Ac)TAGKSLGPR-K(Biotin)

### Pull-down assays

Pull-down experiments were performed as detailed in Wysocka [52] after nuclear extraction with NE-PER commercial kit following manufactureŕs instructions (Thermo Scientific #78833) and run in SDS-PAGE for a short time in order to avoid separation. Protein gels were fixed by soaking for 3 hr in 30% methanol, 2% phosphoric acid. Then the gel was washed with deionized water 3 times, 5 minutes each. After removing the deionized water, the staining solution (0.5 g/L Coomasie blue brilliant R250, 18% methanol, 17% (NH_4_)2SO_4_, 2% phosphoric acid) was added until covering the gel. It was stained for 1 h in gentle shaking. Then the gel was rinsed in deionized water 3 times for 5 minutes each. The background staining was removed with 30% methanol. The gel was stored in deionized water until use. Each lane of the gel was cut into individual slices. Each band was then cut into 1 mm^3^ cube and further treated with three washes of 50 mM NH_4_HCO_3_ in 50% CH_3_CN with 10 min incubations. Each group of gel cubes was then dehydrated in CH_3_CN for 10 min and dried in a Speed Vac. Protein samples were reduced by dithiothreitol (DTT) and alkylated by iodoacetamide [92]. A solution of 10 ng/µL trypsin in 50 mM NH_4_HCO_3_ was used to re-swell the gel pieces completely at 4°C for 30 min, followed by a 37°C digestion overnight. A small amount of 10% formic acid was then added to stop the digestion. The sample was then centrifuged at 2,800 x g, and the supernatant was collected for LC-MS/MS.

### LC-MS/MS Analysis

A fused silica microcapillary LC column (15-cm long x 75-µm inside diameter) packed with Halo C18 reversed-phase resin (2.7 µm particle size, 90 nm pore size, MichromBioresources.) was used with EASY-nLC 1200 system (Thermo Fisher). The nanospray ESI was fitted onto the Thermo Q-Exactive plus mass spectrometer (Thermo Electron, San Jose, CA) that was operated in a Higher-energy C-trap dissociation mode to obtain both MS and tandem MS (MS/MS) spectra. Two µL of tryptic peptide samples were loaded onto the microcapillary column and separated by applying a gradient of 3-40% acetonitrile in 0.1% formic acid at a flow rate of 300 nL/min for 80 min. Mass spectrometry data were acquired in a data-dependent top-10 acquisition mode, which uses a full MS scan from m/z 350-1700 at 70,000 resolution (automatic gain control [AGC] target, 1e6; maximum ion time [max IT], 100 ms; profile mode). Resolution for dd-MS2 spectra was set to 17,500 (AGC target: 1e5) with a maximum ion injection time of 50 ms. The normalized collision energy was 27 eV.

### Protein Identification

Obtained MS spectra were searched against the *T. gondii* ToxoDB-42 _TgondiiME49 protein database using Proteome Discoverer 2.2 (Thermo Electron, San Jose, CA). The search parameters permitted a 10 ppm peptide MS tolerance and a 0.02 Da MS/MS tolerance. Carboxymethylation of cysteines was set as a fixed modification, and oxidation of methionine as a dynamic modification. Up to two missed tryptic peptide cleavages were considered. The proteins for which the False Discovery Rate was less than 1% at the peptide level were included in the following analysis. The raw data of all mass spectra had been submitted to the MassIVE (https://massive.ucsd.edu/; Project accession: MassIVE MSV000091226).

## Author Contributions

L.V. participated in all the experiments, analyses and design of the study and wrote the first draft; D.M. performed Crispr/Cas9 and some of the characterization experiments along with the pull-down experiments with L.V. supervision; C.C. performed the DNA damage experiments and some of the transfections; A.G. was involved in many of the experiments with technical support, and was in charge of RNA and cDNA preparation; R.N. performed RT-PCR experiments and ELISAs with LV supervision; M.C.B. performed Co-IP experiments with L.V. supervison; V.T. perfomed Rop5 and Rop18 antibodies and provided the respective primers for RT-PCR; F.G. synthesized biotynilated peptides for pull down experiments; B.D. performed mass spectrometry analysis; K.K. and W.J.S. participated in design of the study and manuscript final corrections; S.O.A. designed the study and participated in its supervision. All authors have read and agreed to the published version of the manuscript.

## Funding

This work was supported by the Ministerio Nacional de Ciencia y Tecnología (MINCyT): PICT 2015 1288 (S.O.A.), PICT 2018 2434 (L.V.), Consejo Nacional de Investigaciones Científicas y Tecnológicas (CONICET): PIP 11220150100145CO and 11220210100572CO (S.O.A., L.V.) and by National Institute of Health: NIH-NIAID 1R01AI129807 (S.O.A.) and AI152583 (to W.J.S.).

## Institutional Review Board Statement

Not applicable.

## Informed Consent Statement

Not applicable.

## Data Availability Statement

The data that support the findings of this study are available from the corresponding author upon reasonable request.

## Acknowledgments

L. Vanagas (Researcher), D. Muñoz (Fellow), C. Cristaldi (Fellow), D. Ganuza (Technician), V. Turowski (Researcher), and S.O. Angel (Researcher) are members of CONICET. S.O. Angel (Full) and L. Vanagas (Adjunct) are Professors at Universidad Nacional General de San Martin (UNSAM). The Vermont Biomedical Research Network (VBRN) Proteomics Facility is supported through NIH grant P20GM103449 from the INBRE Program of the National Institute of General Medical Science (BD).

## Conflicts of interest

The authors declare that there are no conflict of interest.

## Supplementary figures

**Fig. S1.**
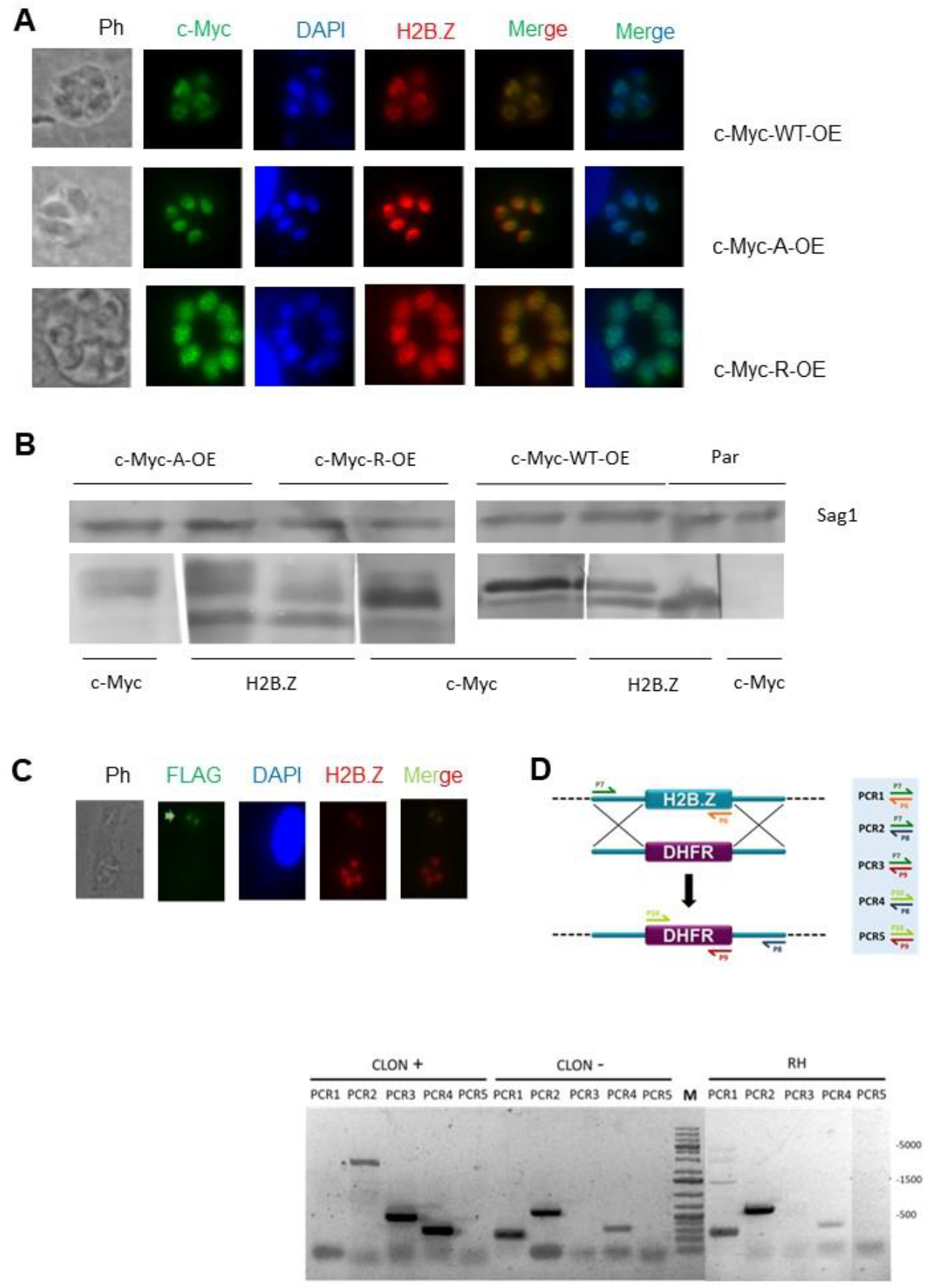
Controls for tachyzoite lines over-expressing H2B.Z WT or mutants and CRISPR/Cas9 deletion of endogenous H2B.Z gene. **A.** Immunofluorescence assay. Htert confluent slides were infected with the different clones obtained and immunofluorescence was performed with anti c-Myc (green) and anti-H2B.Z (red); DAPI was used to stain nuclei. Images show a representative IFA of one clone of each over-expressing line obtained. **B.** Western-blot assay. One clone of each line and the parental were lysed and run in SDS-PAGE for WB assay with anti-H2B.Z, anti-c-Myc and anti-Sag1 (charge control). **C.** Immunofluorescence after transfection showing flag-CAS9 signal in the nucleus of one vacuole (green arrow). **D.** PCR selection of positive clones. The scheme shows the primers designed for PCR detection of positive and negative clones. The image corresponds to a representative PCR where a positive and a negative clone with the parental as control are shown. PCR1-5: primers combinations for PCR shown in the scheme.

**Fig. S2.**
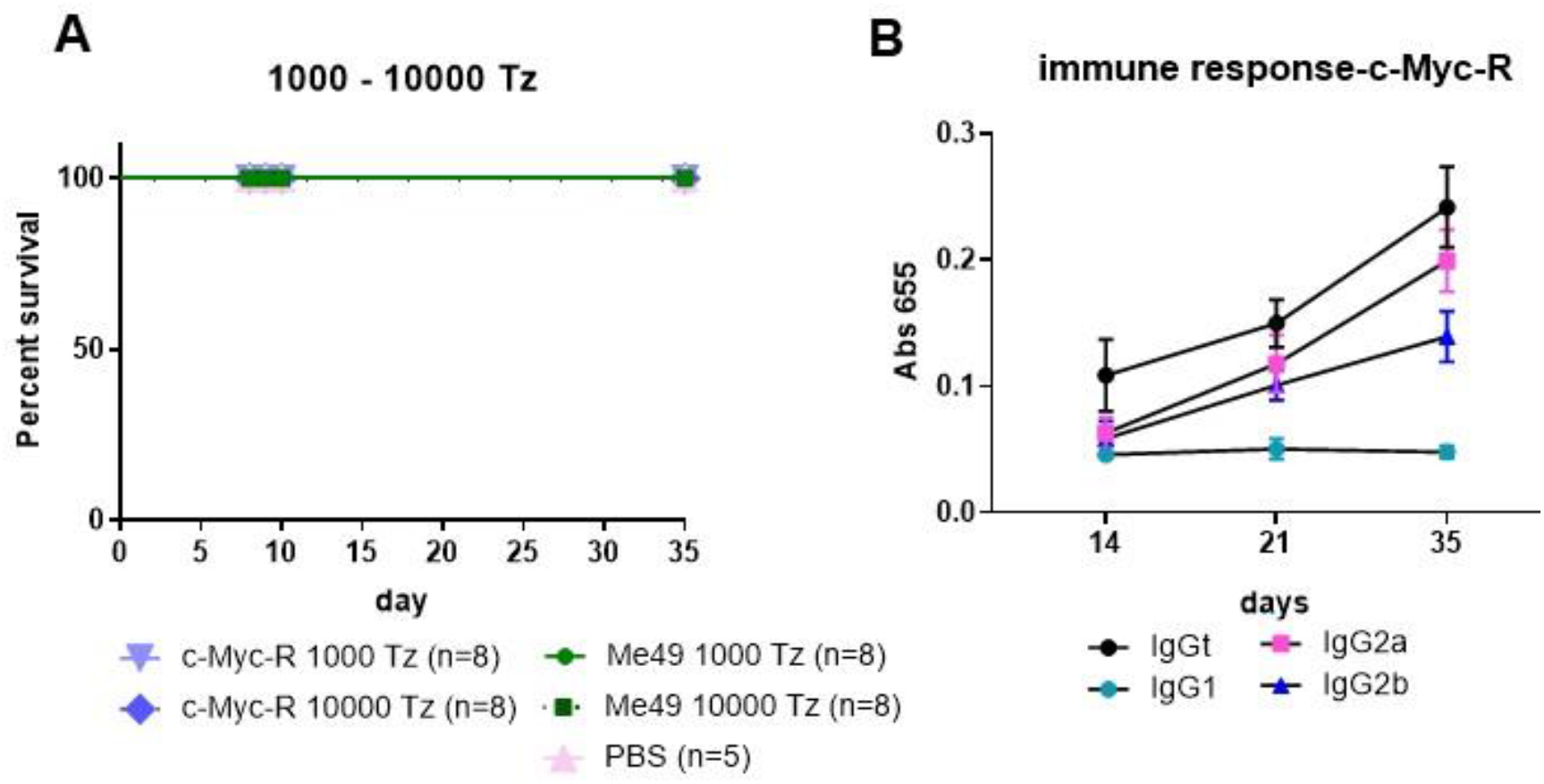
c-Myc-R tachyzoites up to 10,000 are unable to kill mice but immune response is triggered. **A.** Survival assay. 10 mice C57BL/6 were intra-peritoneal infected with either 1000 or 10000 tachyzoites of Me49 or c-Myc-R tachyzoites. Vehicle (PBS) was inoculated in 5 mice as control. The survival curve shown corresponds to one assay, representative of three independent experiments. **B.** Evaluation of immune response by ELISA. Assay shown in Fig. 5C, now showing data for 14, 21 and 35 days samples from mice infeceted with c-Myc-R tachyzoites. Total IgG and subtypes (IgG2a, IgG2b and IgG1) were analyzed.

**Fig. S3.**
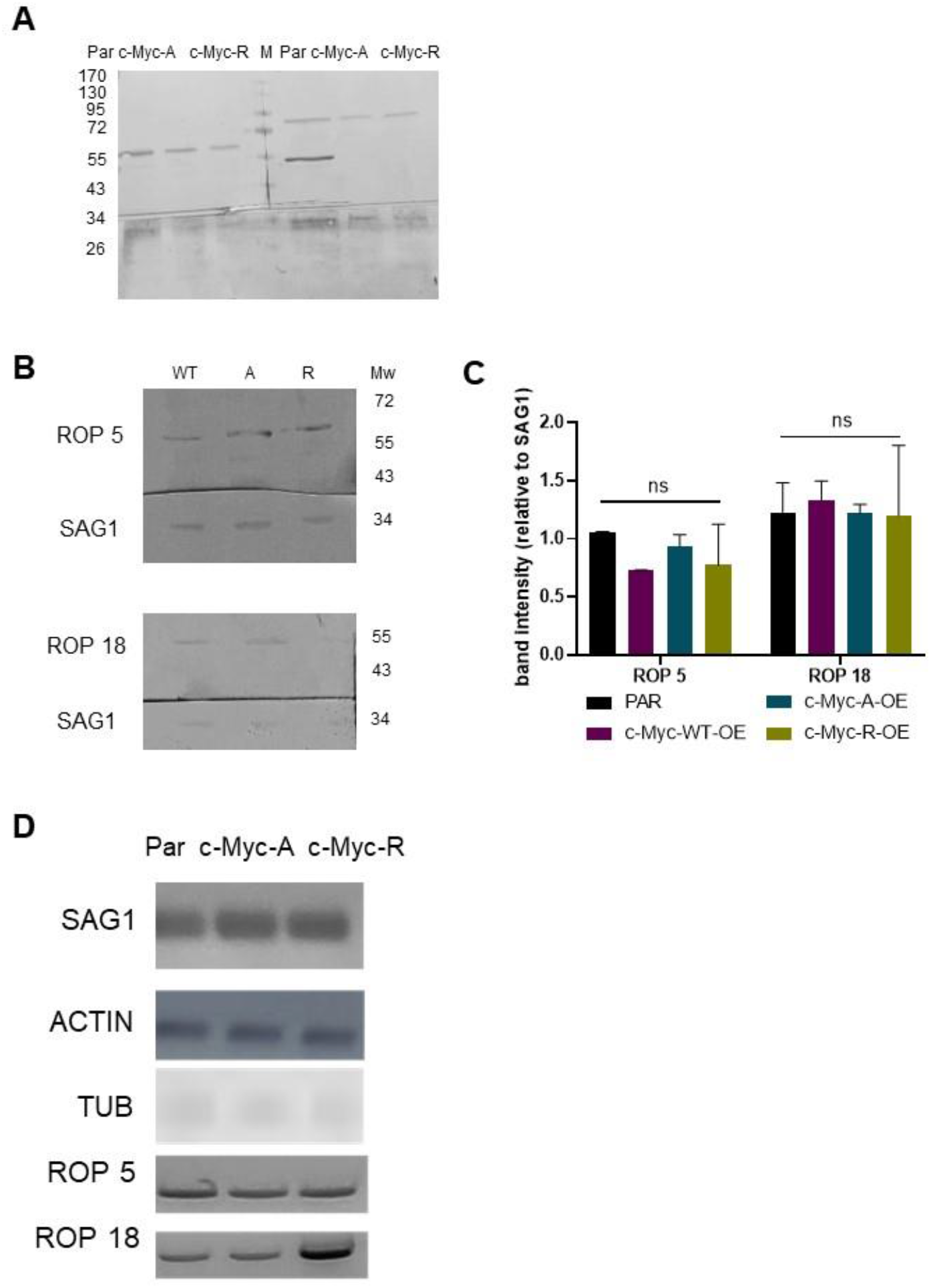
Western blot analysis of ROP proteins. **A.** Complete image of Western blot shown in Fig. 5. **B.** Representative Western-blot of over-expressing lines. Antibodies against Rop5, Rop18 and Sag1 were used. WT: c-Myc-WT-OE; A: c-Myc-A-OE; R: c-Myc-R-OE. The upper bands detected with αRop18 are of higher molecular weight than Rop18 full length protein, being unspecific. However, the intensities are similar to Sag1, being a second charge control. **C.** Quantification of Rop5 and Rop18 bands, relativized to Sag1 band intensity in each lane, in two independent WB experiments with the OE lines, and statistical analysis by GraphPad Prism 8. ns: not significant. **D.** RT-PCR. Tachyzoites of parental (RHΔ*hxgprt*), c-Myc-A and c-Myc-R were collected by triplicate and conserved in TriZol until RNA extraction, and cDNA preparation. PCR was run with primers for the genes indicated and ran in agarose gels. Image is representative of three independent experiments.

**Fig. S4.**
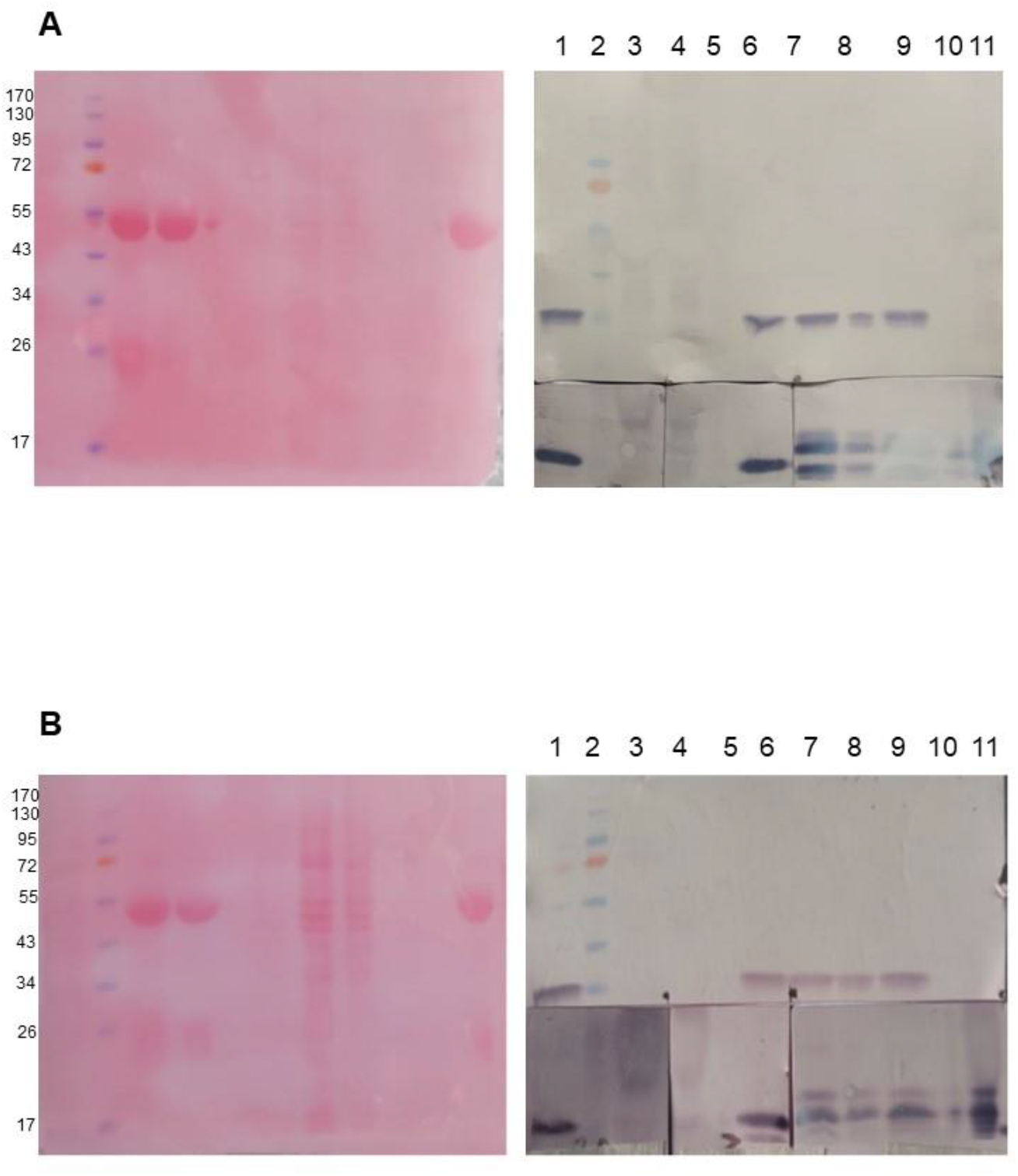
Co-immunoprecipitation assay. **A.** Complete image of the Pounceau (left) and WB (right) shown in Fig. 7 for c-Myc-A. **B.** Complete image of the Pounceau (left) and WB (right) shown in Fig. 7 for c-Myc-R. For both images in lanes 1-11 was run: IN-M-IP-IP-space-IN-IN-T-NU-space-IP. IN: input; IP: immunoprecipitation; T: total; NU: after immunoprecipitation, not bound to agarose beads. MW markers are indicated at the left.

## Notes

https://massive.ucsd.edu/

### Competing Interest Statement

The authors have declared no competing interest.

### Summary of Updates

Grammar corrections and author affiliations updated

## References

[1] Vanagas L, Jeffers V, Bogado SS, Dalmasso MC, Sullivan WJ, Jr., Angel SO. Toxoplasma histone acetylation remodelers as novel drug targets. Expert review of anti-infective therapy. 2012;10:1189–201.

[2] Mouveaux T, Rotili D, Boissavy T, Roger E, Pierrot C, Mai A, et al. A potent HDAC inhibitor blocks Toxoplasma gondii tachyzoite growth and profoundly disrupts parasite gene expression. Int J Antimicrob Agents. 2022;59:106526.

[3] Kim K. The Epigenome, Cell Cycle, and Development in Toxoplasma. Annual review of microbiology. 2018;72:479–99.

[4] Pappas G, Roussos N, Falagas ME. Toxoplasmosis snapshots: global status of Toxoplasma gondii seroprevalence and implications for pregnancy and congenital toxoplasmosis. Int J Parasitol. 2009;39:1385–94.

[5] Attias M, Teixeira DE, Benchimol M, Vommaro RC, Crepaldi PH, De Souza W. The life-cycle of Toxoplasma gondii reviewed using animations. Parasit Vectors. 2020;13:588.

[6] Weiss LM, Kim K. The development and biology of bradyzoites of Toxoplasma gondii. Front Biosci. 2000;5:D391–405.

[7] Luft BJ, Remington JS. Toxoplasmic encephalitis in AIDS. Clin Infect Dis. 1992;15:211–22.

[8] Carlier Y, Truyens C, Deloron P, Peyron F. Congenital parasitic infections: a review. Acta Trop. 2012;121:55–70.

[9] Brynska A, Tomaszewicz-Libudzic E, Wolanczyk T. Obsessive-compulsive disorder and acquired toxoplasmosis in two children. Eur Child Adolesc Psychiatry. 2001;10:200–4.

[10] Miman O, Mutlu EA, Ozcan O, Atambay M, Karlidag R, Unal S. Is there any role of Toxoplasma gondii in the etiology of obsessive-compulsive disorder? Psychiatry Res. 2010;177:263–5.

[11] Yolken RH, Dickerson FB, Fuller Torrey E. Toxoplasma and schizophrenia. Parasite Immunol. 2009;31:706–15.

[12] Vittecoq M, Elguero E, Lafferty KD, Roche B, Brodeur J, Gauthier-Clerc M, et al. Brain cancer mortality rates increase with Toxoplasma gondii seroprevalence in France. Infect Genet Evol. 2012;12:496–8.

[13] de Haan L, Sutterland AL, Schotborgh JV, Schirmbeck F, de Haan L. Association of Toxoplasma gondii Seropositivity With Cognitive Function in Healthy People: A Systematic Review and Meta-analysis. JAMA Psychiatry. 2021;78:1103–12.

[14] Waldman BS, Schwarz D, Wadsworth MH, 2nd, Saeij JP, Shalek AK, Lourido S. Identification of a Master Regulator of Differentiation in Toxoplasma. Cell. 2020;180:359–72 e16.

[15] Jeffers V, Tampaki Z, Kim K, Sullivan WJ, Jr. A latent ability to persist: differentiation in Toxoplasma gondii. Cellular and molecular life sciences : CMLS. 2018;75:2355–73.

[16] Bougdour A, Maubon D, Baldacci P, Ortet P, Bastien O, Bouillon A, et al. Drug inhibition of HDAC3 and epigenetic control of differentiation in Apicomplexa parasites. The Journal of experimental medicine. 2009;206:953–66.

[17] Nardelli SC, Silmon de Monerri NC, Vanagas L, Wang X, Tampaki Z, Sullivan WJ, Jr., et al. Genome-wide localization of histone variants in Toxoplasma gondii implicates variant exchange in stage-specific gene expression. BMC genomics. 2022;23:128.

[18] Santisteban MS, Kalashnikova T, Smith MM. Histone H2A.Z regulats transcription and is partially redundant with nucleosome remodeling complexes. Cell. 2000;103:411–22.

[19] Oberdoerffer P, Miller KM. Histone H2A variants: Diversifying chromatin to ensure genome integrity. Semin Cell Dev Biol. 2022.

[20] Xu Y, Ayrapetov MK, Xu C, Gursoy-Yuzugullu O, Hu Y, Price BD. Histone H2A.Z controls a critical chromatin remodeling step required for DNA double-strand break repair. Mol Cell. 2012;48:723–33.

[21] Fenoy IM, Bogado SS, Contreras SM, Gottifredi V, Angel SO. The Knowns Unknowns: Exploring the Homologous Recombination Repair Pathway in Toxoplasma gondii. Front Microbiol. 2016;7:627.

[22] Dalmasso MC, Echeverria PC, Zappia MP, Hellman U, Dubremetz JF, Angel SO. Toxoplasma gondii has two lineages of histones 2b (H2B) with different expression profiles. Molecular and biochemical parasitology. 2006;148:103–7.

[23] Dalmasso MC, Onyango DO, Naguleswaran A, Sullivan WJ, Jr., Angel SO. Toxoplasma H2A variants reveal novel insights into nucleosome composition and functions for this histone family. J Mol Biol. 2009;392:33–47.

[24] Dalmasso MC, Sullivan WJ, Jr., Angel SO. Canonical and variant histones of protozoan parasites. Frontiers in bioscience. 2011;16:2086–105.

[25] Petter M, Selvarajah SA, Lee CC, Chin WH, Gupta AP, Bozdech Z, et al. H2A.Z and H2B.Z double-variant nucleosomes define intergenic regions and dynamically occupy var gene promoters in the malaria parasite Plasmodium falciparum. Molecular microbiology. 2013;87:1167–82.

[26] Lowell JE, Kaiser F, Janzen CJ, Cross GA. Histone H2AZ dimerizes with a novel variant H2B and is enriched at repetitive DNA in Trypanosoma brucei. J Cell Sci. 2005;118:5721–30.

[27] Sidik SM, Huet D, Ganesan SM, Huynh MH, Wang T, Nasamu AS, et al. A Genome-wide CRISPR Screen in Toxoplasma Identifies Essential Apicomplexan Genes. Cell. 2016;166:1423–35 e12.

[28] Nardelli SC, Che FY, Silmon de Monerri NC, Xiao H, Nieves E, Madrid-Aliste C, et al. The histone code of Toxoplasma gondii comprises conserved and unique posttranslational modifications. mBio. 2013;4:e00922–13.

[29] Giaimo BD, Ferrante F, Herchenrother A, Hake SB, Borggrefe T. The histone variant H2A.Z in gene regulation. Epigenetics & chromatin. 2019;12:37.

[30] Talbert PB, Henikoff S. Histone variants--ancient wrap artists of the epigenome. Nature reviews Molecular cell biology. 2010;11:264–75.

[31] Bruce K, Myers FA, Mantouvalou E, Lefevre P, Greaves I, Bonifer C, et al. The replacement histone H2A.Z in a hyperacetylated form is a feature of active genes in the chicken. Nucleic Acids Res. 2005;33:5633–9.

[32] Binda O, Sevilla A, LeRoy G, Lemischka IR, Garcia BA, Richard S. SETD6 monomethylates H2AZ on lysine 7 and is required for the maintenance of embryonic stem cell self-renewal. Epigenetics. 2013;8:177–83.

[33] Law C, Cheung P. Expression of Non-acetylatable H2A.Z in Myoblast Cells Blocks Myoblast Differentiation through Disruption of MyoD Expression. J Biol Chem. 2015;290:13234–49.

[34] Valdes-Mora F, Song JZ, Statham AL, Strbenac D, Robinson MD, Nair SS, et al. Acetylation of H2A.Z is a key epigenetic modification associated with gene deregulation and epigenetic remodeling in cancer. Genome research. 2012;22:307–21.

[35] Whittle CM, McClinic KN, Ercan S, Zhang X, Green RD, Kelly WG, et al. The genomic distribution and function of histone variant HTZ-1 during C. elegans embryogenesis. PLoS Genet. 2008;4:e1000187.

[36] Updike DL, Mango SE. Temporal regulation of foregut development by HTZ-1/H2A.Z and PHA-4/FoxA. PLoS Genet. 2006;2:e161.

[37] Ridgway P, Brown KD, Rangasamy D, Svensson U, Tremethick DJ. Unique residues on the H2A.Z containing nucleosome surface are important for Xenopus laevis development. J Biol Chem. 2004;279:43815–20.

[38] Myers FA, Chong W, Evans DR, Thorne AW, Crane-Robinson C. Acetylation of histone H2B mirrors that of H4 and H3 at the chicken beta-globin locus but not at housekeeping genes. J Biol Chem. 2003;278:36315–22.

[39] Miao J, Fan Q, Cui L, Li J, Li J, Cui L. The malaria parasite Plasmodium falciparum histones: organization, expression, and acetylation. Gene. 2006;369:53–65.

[40] Trelle MB, Salcedo-Amaya AM, Cohen AM, Stunnenberg HG, Jensen ON. Global histone analysis by mass spectrometry reveals a high content of acetylated lysine residues in the malaria parasite Plasmodium falciparum. Journal of proteome research. 2009;8:3439–50.

[41] Radke JB, Lucas O, De Silva EK, Ma Y, Sullivan WJ, Jr., Weiss LM, et al. ApiAP2 transcription factor restricts development of the Toxoplasma tissue cyst. Proc Natl Acad Sci U S A. 2013;110:6871–6.

[42] Hong DP, Radke JB, White MW. Opposing Transcriptional Mechanisms Regulate Toxoplasma Development. mSphere. 2017;2.

[43] Gursoy-Yuzugullu O, Ayrapetov MK, Price BD. Histone chaperone Anp32e removes H2A.Z from DNA double-strand breaks and promotes nucleosome reorganization and DNA repair. Proc Natl Acad Sci U S A. 2015;112:7507–12.

[44] Morillo-Huesca M, Clemente-Ruiz M, Andujar E, Prado F. The SWR1 histone replacement complex causes genetic instability and genome-wide transcription misregulation in the absence of H2A.Z. PloS one. 2010;5:e12143.

[45] Munera Lopez J, Ganuza A, Bogado SS, Munoz D, Ruiz DM, Sullivan WJ, Jr., et al. Evaluation of ATM Kinase Inhibitor KU-55933 as Potential Anti-Toxoplasma gondii Agent. Frontiers in cellular and infection microbiology. 2019;9:26.

[46] Angel SO, Vanagas L, Ruiz DM, Cristaldi C, Saldarriaga Cartagena AM, Sullivan WJJ. Emerging Therapeutic Targets Against Toxoplasma gondii: Update on DNA Repair Response Inhibitors and Genotoxic Drugs. Front Cell Infect Microbiol. 2020;10.

[47] Behnke MS, Fentress SJ, Mashayekhi M, Li LX, Taylor GA, Sibley LD. The polymorphic pseudokinase ROP5 controls virulence in Toxoplasma gondii by regulating the active kinase ROP18. PLoS pathogens. 2012;8:e1002992.

[48] Etheridge RD, Alaganan A, Tang K, Lou HJ, Turk BE, Sibley LD. The Toxoplasma pseudokinase ROP5 forms complexes with ROP18 and ROP17 kinases that synergize to control acute virulence in mice. Cell host & microbe. 2014;15:537–50.

[49] Reese ML, Zeiner GM, Saeij JP, Boothroyd JC, Boyle JP. Polymorphic family of injected pseudokinases is paramount in Toxoplasma virulence. Proc Natl Acad Sci U S A. 2011;108:9625–30.

[50] Saeij JP, Boyle JP, Coller S, Taylor S, Sibley LD, Brooke-Powell ET, et al. Polymorphic secreted kinases are key virulence factors in toxoplasmosis. Science. 2006;314:1780–3.

[51] Fentress SJ, Behnke MS, Dunay IR, Mashayekhi M, Rommereim LM, Fox BA, et al. Phosphorylation of immunity-related GTPases by a Toxoplasma gondii-secreted kinase promotes macrophage survival and virulence. Cell host & microbe. 2010;8:484–95.

[52] Wysocka J. Identifying novel proteins recognizing histone modifications using peptide pull-down assay. Methods. 2006;40:339–43.

[53] Attar N, Campos OA, Vogelauer M, Cheng C, Xue Y, Schmollinger S, et al. The histone H3-H4 tetramer is a copper reductase enzyme. Science. 2020;369:59–64.

[54] Singh A, Verma S, Modak SB, Chaturvedi MM, Purohit JS. Extra-nuclear histones: origin, significance and perspectives. Mol Cell Biochem. 2022;477:507–24.

[55] Choi YS, Hoon Jeong J, Min HK, Jung HJ, Hwang D, Lee SW, et al. Shot-gun proteomic analysis of mitochondrial D-loop DNA binding proteins: identification of mitochondrial histones. Molecular bioSystems. 2011;7:1523–36.

[56] Zanin MK, Donohue JM, Everitt BA. Evidence that core histone H3 is targeted to the mitochondria in Brassica oleracea. Cell Biol Int. 2010;34:997–1003.

[57] Roson JN, Vitarelli MO, Costa-Silva HM, Pereira KS, Pires DDS, Lopes LS, et al. H2B.V demarcates divergent strand-switch regions, some tDNA loci, and genome compartments in Trypanosoma cruzi and affects parasite differentiation and host cell invasion. PLoS pathogens. 2022;18:e1009694.

[58] Croken MM, Nardelli SC, Kim K. Chromatin modifications, epigenetics, and how protozoan parasites regulate their lives. Trends Parasitol. 2012;28:202–13.

[59] Croken MM, Qiu W, White MW, Kim K. Gene Set Enrichment Analysis (GSEA) of Toxoplasma gondii expression datasets links cell cycle progression and the bradyzoite developmental program. BMC genomics. 2014;15:515.

[60] Wang J, Dixon SE, Ting LM, Liu TK, Jeffers V, Croken MM, et al. Lysine acetyltransferase GCN5b interacts with AP2 factors and is required for Toxoplasma gondii proliferation. PLoS pathogens. 2014;10:e1003830.

[61] Jeffers V, Gao H, Checkley LA, Liu Y, Ferdig MT, Sullivan WJ, Jr. Garcinol Inhibits GCN5-Mediated Lysine Acetyltransferase Activity and Prevents Replication of the Parasite Toxoplasma gondii. Antimicrobial agents and chemotherapy. 2016;60:2164–70.

[62] Huang S, Holmes MJ, Radke JB, Hong DP, Liu TK, White MW, et al. Toxoplasma gondii AP2IX-4 Regulates Gene Expression during Bradyzoite Development. mSphere. 2017;2.

[63] Khelifa AS, Guillen Sanchez C, Lesage KM, Huot L, Mouveaux T, Pericard P, et al. TgAP2IX-5 is a key transcriptional regulator of the asexual cell cycle division in Toxoplasma gondii. Nature communications. 2021;12:116.

[64] Sindikubwabo F, Ding S, Hussain T, Ortet P, Barakat M, Baumgarten S, et al. Modifications at K31 on the lateral surface of histone H4 contribute to genome structure and expression in apicomplexan parasites. eLife. 2017;6.

[65] Gissot M, Kelly KA, Ajioka JW, Greally JM, Kim K. Epigenomic modifications predict active promoters and gene structure in Toxoplasma gondii. PLoS pathogens. 2007;3:e77.

[66] Ren Q, Gorovsky MA. Histone H2A.Z acetylation modulates an essential charge patch. Mol Cell. 2001;7:1329–35.

[67] Ren Q, Gorovsky MA. The nonessential H2A N-terminal tail can function as an essential charge patch on the H2A.Z variant N-terminal tail. Mol Cell Biol. 2003;23:2778–89.

[68] Macadangdang BR, Oberai A, Spektor T, Campos OA, Sheng F, Carey MF, et al. Evolution of histone 2A for chromatin compaction in eukaryotes. eLife. 2014;3.

[69] Severo V, Souza R, Vitorino F, Cunha J, Avila A, Arrizabalaga G, et al. Previously Unidentified Histone H1-Like Protein Is Involved in Cell Division and Ribosome Biosynthesis in Toxoplasma gondii. mSphere. 2022;7:e0040322.

[70] Sarah Sokol Borrelli SMR, Katherine G. Sharp, Leah F. Cabo, Hisham S. Alrubaye, Bruno Martorelli Di Genova and Jon P. Boyle. A transcriptional network required for Toxoplasma gondii tissue cyst formation is dispensable for long-term persistence. bioRxiv. 2022.

[71] M. Haley Licon CJG, Sundeep Chakladar, Lindsey Shallberg, Benjamin S. Waldman, Christopher A. Hunter, Sebastian Lourido. A positive feedback loop controls Toxoplasma chronic differentiation. Biorxvi. 2022.

[72] Colino-Sanguino Y, Cornett EM, Moulder D, Smith GC, Hrit J, Cordeiro-Spinetti E, et al. A Read/Write Mechanism Connects p300 Bromodomain Function to H2A.Z Acetylation. iScience. 2019;21:773–88.

[73] Afroz T, Cienikova Z, Clery A, Allain FHT. One, Two, Three, Four! How Multiple RRMs Read the Genome Sequence. Methods in enzymology. 2015;558:235–78.

[74] Malumbres M. Cyclin-dependent kinases. Genome biology. 2014;15:122.

[75] Jin C, Zang C, Wei G, Cui K, Peng W, Zhao K, et al. H3.3/H2A.Z double variant-containing nucleosomes mark ‘nucleosome-free regions’ of active promoters and other regulatory regions. Nature genetics. 2009;41:941–5.

[76] Rahal R, Amon A. Mitotic CDKs control the metaphase-anaphase transition and trigger spindle elongation. Genes & development. 2008;22:1534–48.

[77] Lukasik P, Zaluski M, Gutowska I. Cyclin-Dependent Kinases (CDK) and Their Role in Diseases Development-Review. International journal of molecular sciences. 2021;22.

[78] Ors A, Papin C, Favier B, Roulland Y, Dalkara D, Ozturk M, et al. Histone H3.3 regulates mitotic progression in mouse embryonic fibroblasts. Biochemistry and cell biology = Biochimie et biologie cellulaire. 2017;95:491–9.

[79] Page MJ, Di Cera E. Evolution of peptidase diversity. J Biol Chem. 2008;283:30010–4.

[80] Harvey SH, Krien MJ, O’Connell MJ. Structural maintenance of chromosomes (SMC) proteins, a family of conserved ATPases. Genome biology. 2002;3:REVIEWS3003.

[81] Laflamme G, Tremblay-Boudreault T, Roy MA, Andersen P, Bonneil E, Atchia K, et al. Structural maintenance of chromosome (SMC) proteins link microtubule stability to genome integrity. J Biol Chem. 2014;289:27418–31.

[82] Serrano D, Cordero G, Kawamura R, Sverzhinsky A, Sarker M, Roy S, et al. The Smc5/6 Core Complex Is a Structure-Specific DNA Binding and Compacting Machine. Mol Cell. 2020;80:1025–38 e5.

[83] Borgogno MV, Monti MR, Zhao W, Sung P, Argarana CE, Pezza RJ. Tolerance of DNA Mismatches in Dmc1 Recombinase-mediated DNA Strand Exchange. J Biol Chem. 2016;291:4928–38.

[84] Denkers EY, Gazzinelli RT. Regulation and function of T-cell-mediated immunity during Toxoplasma gondii infection. Clin Microbiol Rev. 1998;11:569–88.

[85] Blader IJ, Saeij JP. Communication between Toxoplasma gondii and its host: impact on parasite growth, development, immune evasion, and virulence. APMIS. 2009;117:458–76.

[86] Elsaid MM, Vitor RW, Frezard FJ, Martins MS. Protection against toxoplasmosis in mice immunized with different antigens of Toxoplasma gondii incorporated into liposomes. Mem Inst Oswaldo Cruz. 1999;94:485–90.

[87] Sanchez-Lopez EF, Corigliano MG, Oliferuk S, Ramos-Duarte VA, Rivera M, Mendoza-Morales LF, et al. Oral Immunization With a Plant HSP90-SAG1 Fusion Protein Produced in Tobacco Elicits Strong Immune Responses and Reduces Cyst Number and Clinical Signs of Toxoplasmosis in Mice. Front Plant Sci. 2021;12:726910.

[88] Buxton D, Innes EA. A commercial vaccine for ovine toxoplasmosis. Parasitology. 1995;110 Suppl:S11-6.

[89] Wu M, Liu S, Chen Y, Liu D, An R, Cai H, et al. Live-attenuated ME49Deltacdpk3 strain of Toxoplasma gondii protects against acute and chronic toxoplasmosis. NPJ Vaccines. 2022;7:98.

[90] Echeverria PC, Matrajt M, Harb OS, Zappia MP, Costas MA, Roos DS, et al. Toxoplasma gondii Hsp90 is a potential drug target whose expression and subcellular localization are developmentally regulated. J Mol Biol. 2005;350:723–34.

[91] Guzman F, Gauna A, Roman T, Luna O, Alvarez C, Pareja-Barrueto C, et al. Tea Bags for Fmoc Solid-Phase Peptide Synthesis: An Example of Circular Economy. Molecules. 2021;26.

[92] Spiess PC, Deng B, Hondal RJ, Matthews DE, van der Vliet A. Proteomic profiling of acrolein adducts in human lung epithelial cells. J Proteomics. 2011;74:2380–94.

